# Characterization of a C9orf72 Knockout *Danio rerio* model for ALS and cross-species validation of potential therapeutics screened in *Caenorhabditis elegans*

**DOI:** 10.1101/2025.03.24.644908

**Authors:** Alexandre Emond, Carl Laflamme, Martine Therrien, Meijiang Liao, Claudia Maios, Audrey Labarre, Pierre Drapeau, J. Alex Parker

**Affiliations:** CHUM Research Center, Montreal, Canada; Department of Neuroscience, University of Montreal, Montreal, Quebec, Canada; Montreal Neurological Institute, McGill University, Montreal, Quebec, Canada; Center for Neuroscience, University of California, Davis, United States of America

## Abstract

Intronic hexanucleotide repeat expansions in the *C9orf72* gene represent the most common genetic cause of the neurodegenerative diseases amyotrophic lateral sclerosis (ALS) and frontotemporal dementia. This expansion decreases *C9orf72* expression in affected patients, indicating that loss of *C9orf72* function (LOF) acts as a pathogenic mechanism. Several models using *Danio rerio* (zebrafish) for *C9orf72* depletion have been developed to explore disease mechanisms and the consequences of *C9orf72* LOF. However, inconsistencies exist in reported phenotypes, and many have yet to be validated in stable germline ablation models. To address this, we created a zebrafish *C9orf72* knockout model using CRISPR/Cas9. The *C9orf72* LOF model demonstrates, in a generally dose-dependent manner, increased larval mortality, persistent growth reduction, and motor deficits. Additionally, homozygous *C9orf72* LOF larvae exhibited mild overbranching of spinal motoneurons. To identify potential therapeutic compounds, we performed a screen on an established *Caenorhabditis elegans* (*C. elegans*) *C9orf72* homologue (*alfa-1*) LOF model, identifying 12 compounds that enhanced motility, reduced neurodegeneration, and alleviated paralysis phenotypes. Motivated by the shared motor phenotype, 2 of those compounds were tested in our zebrafish *C9orf72* LOF model. Pizotifen malate was found to significantly improve motor deficits in *C9orf72* LOF zebrafish larvae. We introduce a novel zebrafish *C9orf72* knockout model that exhibits phenotypic differences from depletion models, providing a valuable tool for *in vivo C9orf72* research and ALS therapeutic validation. Furthermore, we identify pizotifen malate as a promising compound for further preclinical evaluation.

**Author Summary:** Amyotrophic lateral sclerosis (ALS) is a fatal neurodegenerative disorder characterized by the progressive loss of motor neurons, with no curative treatments currently available. The most common genetic cause is a hexanucleotide repeat expansion in the *C9orf72* gene, which reduces its expression and implicates loss-of-function (LOF) as a disease mechanism. However, the complete functions of C9orf72 and its role in ALS remain unclear. Zebrafish models with indirect partial reduction of *C9orf72* expression have shown promise in recapitulating key aspects of ALS, but inconsistencies have been observed across these models. To address these challenges, we developed a stable genetic *C9orf72* LOF zebrafish model to study the effects of its LOF, validate previous findings, and test potential ALS therapeutics. Our model displays swimming activity deficits, reduced growth, increased mortality, and mild spinal motor neuron abnormalities. We demonstrated that pizotifen malate significantly improved motor function in both our model and a similar well-established worm model. These results underscore the differences between indirect depletion and direct genetic LOF models while identifying pizotifen malate as a promising candidate for preclinical testing. This zebrafish model serves as a valuable tool for understanding *C9orf72*-associated ALS mechanisms and advancing therapeutic development.

## Introduction

Amyotrophic lateral sclerosis (ALS) is a devastating neuromuscular disorder characterized by progressive degeneration of motor neurons (MNs). Approved treatments provide only limited benefits, and death typically occurs within 2–3 years post-diagnosis [1]. A GGGGCC hexanucleotide repeat expansion in the first intron of *C9orf72* is the most common genetic cause of ALS (*C9*-ALS) and frontotemporal dementia [2–4]. This expansion is proposed to contribute to ALS pathogenesis through three non-mutually exclusive mechanisms: (1) RNA toxicity via sense and antisense foci, (2) protein toxicity through dipeptide repeat proteins generated by repeat-associated non-ATG translation, and (3) *C9orf72* loss-of-function (LOF), as evidenced by reduced mRNA and protein levels in patient-derived tissues [2,5,6]. Mammalian gain-of-function (GOF) models often fail to fully replicate C9orf72-ALS histopathological and behavioral phenotypes [7]. LOF remains comparatively understudied, and the full biological function of C9orf72 is still unclear [5].

LOF animal models have provided critical insights into *C9orf72* function and ALS pathogenesis. However, *C9orf72* knockout (KO) mouse models fail to fully recapitulate *C9*-ALS pathology, lacking MN loss, exhibiting broad immune dysregulation, and presenting inconsistent motor phenotypes [8–11]. Whether mice have intrinsic resilience to *C9orf72* LOF-induced motor deficits and MN degeneration remains unclear. This highlights the need for alternative vertebrate systems to investigate *C9orf72* LOF and model ALS pathogenesis.

Zebrafish (*Danio rerio*) are a well-established vertebrate model for neurological diseases, sharing key neuroanatomical, neurochemical, and genetic similarities with humans [12]. Many zebrafish models exhibit neurodegenerative markers, including neuronal loss and gliosis, even at early larval stages [13–16]. Zebrafish possess a single *C9orf72* ortholog with 85% sequence homology to the human protein. The first *C9orf72* knockdown (KD) model, generated using morpholinos, revealed spinal MNs axonal overbranching, shortening, and motor deficits [17]. A subsequent microRNA-based KD model demonstrated early mortality, morphological defects, synaptic dysfunction at the neuromuscular junction (NMJ), motor deficits, MNs degeneration, and muscle atrophy [18]. However, a recent CRISPR/Cas9 *C9orf72* KO model exhibited retinal disruption, neuroinflammation, and neurodegeneration in aged mutants, but no spinal MN loss or NMJ defects [19]. Validation of *C9orf72* KD models ALS-related phenotypes in stable germline KO lines is crucial to define zebrafish potential as an alternative vertebrae system for the study of *C9orf72* LOF and ALS pathogenesis.

Non-vertebrate animal models offer advantages for high-throughput compound screening, with promising hits able to be subsequently validated in higher-order systems [20]. *C. elegans* is a well-established model for neurological diseases, with a simple, well-characterized nervous system that shares conserved neuronal subtypes and molecular pathways with humans [21,22]. The sole *C. elegans* ortholog of *C9orf72*, *alfa-1*, shares 59% sequence similarity with its human counterpart. The *alfa-1* KO *C. elegans* mutant line *alfa-1*(ok3062) exhibits age-dependent paralysis and GABAergic MNs degeneration [23]. A robust motor phenotype would enable high-throughput phenotypic screening for *C9*-ALS therapeutics in this line [24–27].

In this study, we created a stable *C9orf72* KO zebrafish model using CRISPR/Cas9 to investigate ALS-related phenotypes, resolve discrepancies with existing KD models, and validate potential *C9*-ALS therapeutics. Homozygous *C9orf72^-/-^* larvae displayed reduced survival, persistent growth deficits, motor impairments, and mild disruption of spinal MN axonal branching and neuromuscular junction (NMJ) postsynaptic organization. In contrast, the heterozygous *C9orf72^-/+^* specimens showed milder mortality, smaller size, and motor deficits. Phenotypes for both genotypes were less severe than those reported for the KD models. Due to the zebrafish’s lesser suitability for high-throughput drug screening, *C. elegans* was used to identify potential *C9*-ALS therapeutics, with the best candidates subsequently validated in our *C9orf72* LOF model. We demonstrated that the *alfa-1(ok3062)* mutant exhibits rapid-onset motor deficits in liquid culture, allowing for the identification of 80 compounds that significantly improved swimming activity in young adult mutant worms. Twelve of these compounds further alleviated age-dependent paralysis and neurodegeneration in *alfa-1(ok3062)* mutants on solid media. By utilizing the motor phenotype of our *C9orf72* KO zebrafish model, we validated pizotifen malate’s (PM) effectiveness in mitigating *C9orf72* LOF-associated motor deficits. Our findings indicate that *C9orf72* LOF induces ALS-related phenotypes in zebrafish larvae, with dose-dependent severity for most phenotypes. However, we did not observe clear signs of spinal MN degeneration at the larval stage. These observations support the notion that stable genetic mutants may show significant differences in phenotypic manifestation and severity compared to RNA interference (RNAi)-based KD models.

## Results

### Generation and validation of a *C9orf72* zebrafish knockout model

Zebrafish have a single conserved ortholog of *C9orf72*, located on chromosome 13 [18]. Using CRISPR/Cas9-mediated genome editing, we generated a *C9orf72* KO line harboring two distinct deletions (Δ4 and Δ7) within the first exon, resulting in a combined deletion of 11 base pairs compared to wild-type (*C9orf72^+/+^*) specimens (**Fig. 1A-B**). These deletions are predicted to cause a frameshift mutation, resulting in premature translation termination and a truncated protein lacking more than 85% of the zebrafish C9orf72 sequence (**Fig. 1C**). To minimize off-target effects, heterozygous *C9orf72^+/-^* F1 individuals were outcrossed with wild-type larvae, and F2 incrossing produced homozygous *C9orf72^-/-^* animals. To increase our confidence in the absence of heritable off-target mutations, we sequenced *C9orf72^-/-^* specimens at the three genomic regions most susceptible to off-target cleavage, as predicted by the CRISPRoff tool [28]. These regions included intron 2 of *Pou6f1*, intron 3 of *si:dkey-277i15.2*, and exon 11 of *ch25hl3*. Sequencing revealed no evidence of off-target cleavage, as all regions were identical to those in *C9orf72*^+/+^ controls (**S1 Fig)**

**Fig. 1.**
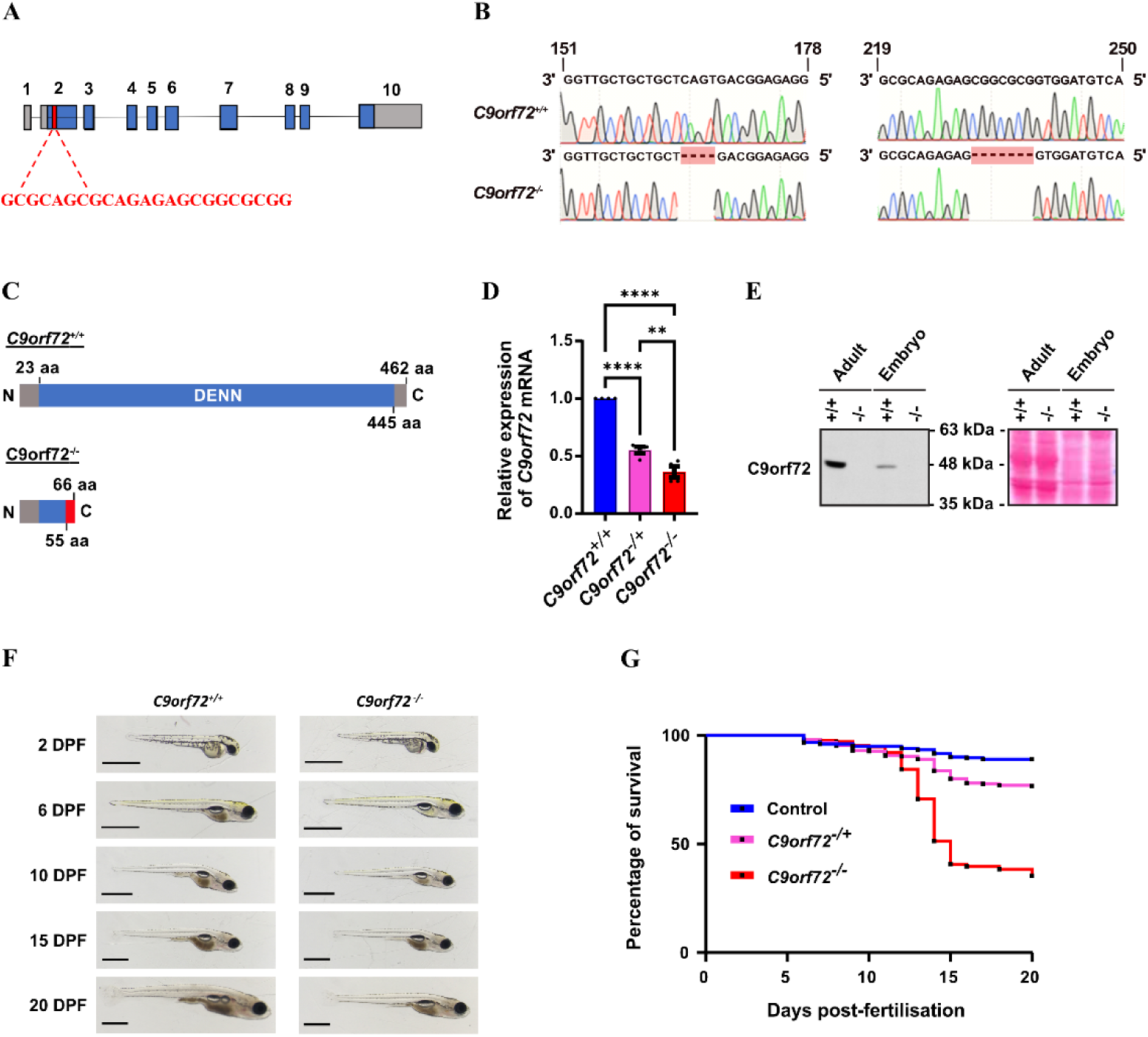
Generation and validation of a stable zebrafish *C9orf72* knockout line. **(A)** Schematic representation of the CRISPR/Cas9-mediated knockout of the endogenous *C9orf72* gene. A guide RNA was designed to target the start codon region in the second exon of C9orf72, inducing indels. **(B)** Sequencing of the *C9orf72* knockout line revealed two distinct deletions (Δ4 and Δ7) at different sites within the *C9orf72* sequence, resulting in a total deletion of 11 base pairs that was absent in *C9orf72*^+/+^ wild-type specimens. **(C)** Schematic representation of the predicted truncated C9orf72 protein resulting from the frameshift caused by the Δ4 and Δ7 deletions. Amino acids (aa). **(D)** Quantitative analysis of *C9orf72* mRNA expression at 2 days post-fertilization (dpf) demonstrated significantly reduced transcript levels in *C9orf72*^+/-^ and *C9orf72^-/-^* compared to *C9orf72*^+/+^ controls, with *C9orf72^-/-^* showing significantly lower levels than *C9orf72*^+/-^. Data are presented as mean ± SEM, normalized to *Polr2d* mRNA levels. Ordinary one-way ANOVA with Holm-Šídák’s multiple comparisons post-hoc test; **** *p < 0.0001*, *\*\* p ≤ 0.01*, * *p ≤ 0.05*; N = 4 experimental repeats, n = 3 technical replicates. **(E)** Immunoblotting and Ponceau staining confirmed the absence of C9orf72 protein in *C9orf72^-/-^ (*-/-) adults and 2 dpf embryos. *C9orf72*^+/+^ (+/+) specimens served as controls for the presence of the protein. **(F)** Comparative morphological analysis revealed no apparent defects in *C9orf72^-/-^* compared to *C9orf72*^+/+^ controls. **(G)** Kaplan–Meier survival analysis up to 20 dpf showed significantly reduced survival in *C9orf72*^+/-^ and *C9orf72^-/-^* compared to *C9orf72*^+/+^ controls after 10 dpf, with *C9orf72^-/-^* exhibiting significantly lower survival than *C9orf72*^+/-^ after 13 dpf. Log-rank (Mantel-Cox) test; N = 3 experimental repeats, n = 300 total fish per genotype. Scale bars = 1 mm.

To validate the LOF in our *C9orf72* KO line, we performed reverse transcription-quantitative polymerase chain reaction (RT-qPCR) using zebrafish-specific TaqMan probes targeting *C9orf72* mRNA. We observed a significant reduction in *C9orf72* mRNA levels in 2 days post fertilization (dpf) larvae: 57.8 ± 6.7% in *C9orf72^-/+^* and 21.2% ± 2.1% in *C9orf72^-/-^* compared to *C9orf72^+/+^* wild-type controls (set at 100%; *p = 0.0041 and* p *< 0.0001,* respectively) (**Fig. 1D**). These results suggest that the frameshift mutation likely triggers nonsense-mediated decay [29], confirming that the introduced indels effectively induce *C9orf72* LOF. To further validate the *C9orf72* KO line, we performed Western blot analysis. As the only previously reported antibody for zebrafish C9orf72 protein (Novus Npb2-15656) [18] had been discontinued, we screened eight antibodies known to recognize mammalian C9orf72 [30]. Of these, only Abcam ab221137 and GeneTex GTX634482 detected zebrafish C9orf72, with Abcam ab221137 demonstrating superior specificity (**S2 Fig.)**. Using ab221137, we confirmed the complete absence of C9orf72 protein in both adult and 2 dpf *C9orf72*^-/-^ specimens (**Fig. 1E**). Together, these findings demonstrate that our genetic approach successfully abolishes C9orf72 protein production *in vivo*. Thus, this *C9orf72* KO line can be used as a tool to investigate the role of *C9orf72* LOF in ALS and its fundamental biological functions.

### Phenotypic characterization of *C9orf72* knockout zebrafish larvae

No overt morphological abnormalities were observed in *C9orf72^-/+^* or *C9orf72^-/-^* specimens during embryonic development (0–3 dpf) or up to 20 dpf (**Fig. 1F**). However, the depletion of C9orf72 resulted in a significant (*p < 0.0001*) decrease in survival from 10 dpf onward, compared to wild-type controls. Survival rates at 20 dpf were 89 ± 1.8%, 76.7 ± 2.4%, and 35 ± 2.8% for *C9orf72^+/+^*, C*9orf72^-/+,^* and *C9orf72^-/-^* specimens, respectively (**Fig. 1G**). Additionally, we observed a persistent and statistically significant reduction in body length (BL) from 2 to 20 dpf in both *C9orf72^-/+^* and *C9orf72^-/-^* specimens, compared to *C9orf72^+/+^* larvae (**Fig. 2A, B**). The magnitude of the growth deficit was dose-dependent, with *C9orf72^-/-^* specimens exhibiting a more pronounced and consistent reduction across all time points (**Fig. 2C-H, S1 Table**).

**Fig 2.**
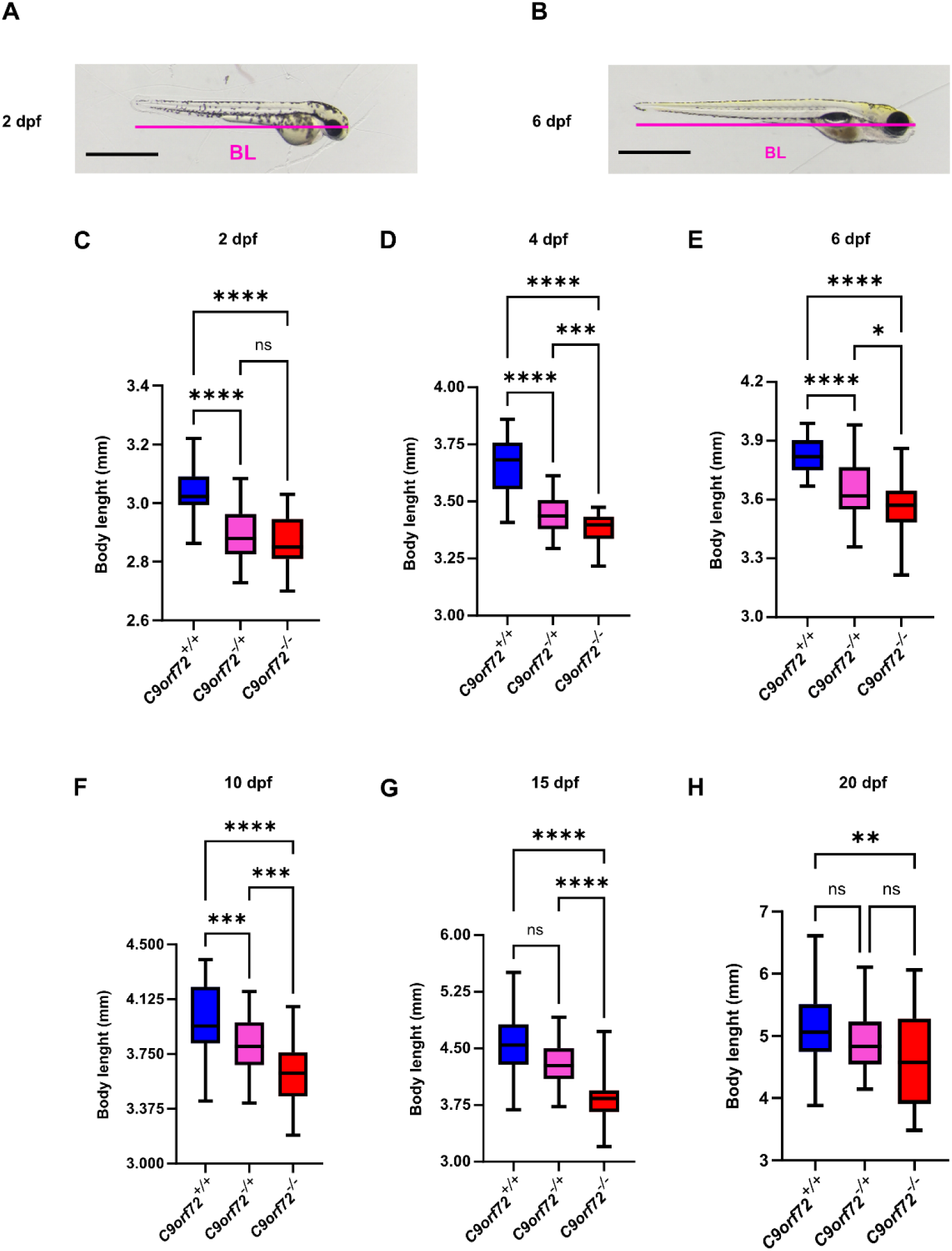
Knockout of C9orf72 results in a persistent reduction in size during late embryonic and larval stages. **(A)** Schematic representation of zebrafish larvae body length (BL) measurement at 2 and 4 days post-fertilization (dpf). **(B)** Schematic representation of BL measurement from 6 dpf onward. **(C-H)** Standard length analysis at 2, 4, 6, 10, 15, and 20 dpf of *C9orf72^+/+^*, *C9orf72^-/+,^* and *C9orf72^-/-^* specimens. **(C)** Significant size reduction observed for *C9orf72^-/+^* and *C9orf72^-/-^* specimens compared to *C9orf72^+/+^* controls at 2 dpf. **(D-F) The** *C9orf72^-/+^* and *C9orf72^-/-^* specimens remain significantly smaller than *C9orf72^+/+^* controls at 4, 6, and 10 dpf, with *C9orf72^-/-^* specimens also significantly shorter than *C9orf72^-/+^* at these time points. **(G)** At 15 dpf, *C9orf72^-/-^* specimens are significantly smaller than both *C9orf72^+/+^* and *C9orf72-/+*. **(H)** At 20 dpf, *C9orf72^-/-^* specimens stay significantly smaller than *C9orf72^+/+^* controls. Statistical tests: ordinary one-way ANOVA with Tukey’s multiple comparisons post-hoc test (**C, F**); Welch and Brown-Forsythe ANOVA with Dunnett’s T3 multiple comparisons post-hoc test (**D, E**); Kruskal-Wallis test with Dunn’s multiple comparisons post-hoc test (**G, H**). **** *p < 0.0001*, *** *p ≤ 0.001*, ** p *< 0.01*, * *p ≤ 0.05,* and NS *p > 0.05*. Boxplot extremities indicate maximum and minimum values; box limits show the interquartile range (central 50%), and the central line marks the median value. n = 35–45 individual larvae per genotype. Scale bars = 1 mm.

To assess the impact of *C9orf72* LOF on motor function, we analyzed zebrafish larval behavior at multiple developmental stages (6, 8, 10, 12, 15, and 20 dpf). As zebrafish larvae exhibit highly stereotyped swimming behaviors even under alternating light/dark conditions [31], we employed a 120-minute alternating dark-light paradigm to assess locomotor activity (**Fig. 3A, B**). This paradigm integrates both autonomous light/dark swimming behavior and acute responses to illumination changes. As larval size influences swimming performance, total locomotor activity was normalized to mean (BL) for each genotype at the corresponding age. Our analysis revealed a significant and persistent locomotor deficit in *C9orf72^-/+^* and *C9orf72^-/-^* larvae compared to *C9orf72^+/+^* controls at 6, 8, 10, 15, and 20 dpf (**Fig. 3C-H**). The locomotor deficit was dose-dependent, with *C9orf72^-/-^* larvae exhibiting a more severe impairment compared to *C9orf72^-/+^* larvae until 15 dpf. Notably, the most pronounced motor differences among genotypes, especially between *C9orf72^-/+^* and *C9orf72^-/-^* larvae, occurred during the early recovery phase following the dark-to-light transition (**Fig. 3A, S3 Fig.**). The *C9orf72* LOF-dependent locomotor deficit was also evident under a 2-phase paradigm (20-minute dark / 60-minute light; (**S4 Fig.**) and persisted even when only considering the stable spontaneous swimming period (SSAP) (**S5 Fig.**). The SSAP was defined as the 40–80 min window, following the 20–40 min adaptation phase after the light transition.

**Fig 3.**
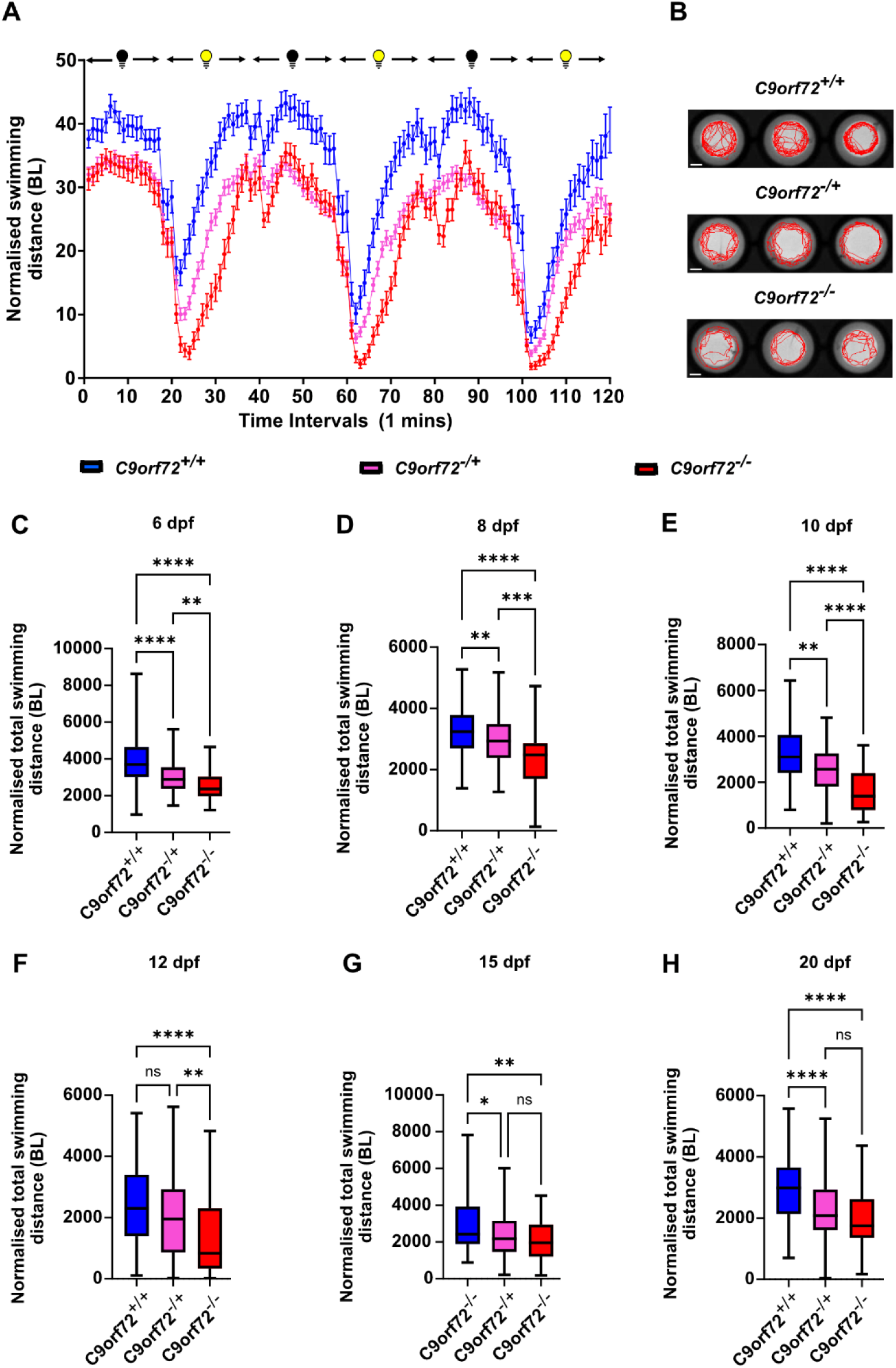
The knockout of *C9orf72* leads to a sustained decrease in swimming activity in larvae. **(A)** Typical swimming distance normalized by mean body length (BL) per minute pattern observed during a 120-minute alternating dark-light paradigm, consisting of 20-minute periods, for *C9orf72^+/+^*, *C9orf72^-/+,^* and *C9orf72^-/-^* 6 days post-fertilization (dpf) larvae. Data are presented as mean ± SEM. (**B)** Representative 60-second swimming tracks of *C9orf72^+/+^*, *C9orf72^-/+,^* and *C9orf72^-/-^* 6 dpf zebrafish larvae subjected to the phasic program. (**C-H)** Quantitative analysis of total swimming distance normalized by corresponding genotype mean BL at 6, 8, 10, 12, 15, and 20 dpf after 120 minutes of exposure to our phasic program. (**C-E**) A significant deficit in swimming activity is observed for C9orf72^-/+^ and *C9orf72^-/-^* compared to *C9orf72^+/+^* controls, and for *C9orf72^-/-^* compared to *C9orf72^+/-^* specimens at 6, 8, and 10 dpf. (**F**) C9orf72^-/-^ specimens show a significant deficit in normalized swimming activity compared to *C9orf72^+/+^* and *C9orf72^-/+^* larvae at 12 dpf. (**G-H**) C9orf72^-/-^ and C9orf72^-/+^ specimens exhibit a significant reduction in normalized swimming activity compared to *C9orf72^+/+^* controls at 15 and 20 dpf. Statistical tests: Kruskal-Wallis test with Dunn’s multiple comparisons post-hoc test (**C, E, F, G, H**), Brown-Forsythe ANOVA with Dunnett’s T3 multiple comparisons post-hoc test (**D**). **** *p < 0.0001*, ** *p ≤ 0.01*, * *p ≤ 0.05,* and NS *p > 0.05*. Boxplot extremities indicate maximum and minimum values, while box limits represent the range of the central 50% of the data, and the central line marks the median value. N = 3, n = 72 for each genotype except for *C9orf72^-/+,^* where n = 144. N represents the number of experimental repeats from different clutches, and n signifies the total number of larvae per genotype considered for the assay. Scale bars = 0.2 cm.

To identify potential neuroanatomical causes underlying the motor deficits observed between 6 and 20 dpf in *C9orf72* KO larvae, we examined the morphological features of spinal primary motoneurons (PMNs). In zebrafish, PMNs are large, segmentally organized MNs located in the spinal cord, projecting ventrally and laterally to innervate the ventral trunk musculature within each spinal hemisegment [32]. To visualize PMNs *in vivo*, we crossed *C9orf72^-/-^* specimens with transgenic *Hb9:GFP* zebrafish (*Tg(mnx1:GFP)*), generating a stable *Hb9C9orf72^-/-^* line. The *Mnx1* promoter drives green fluorescent protein (GFP) expression primarily in motoneurons, allowing us to assess the morphology of spinal PMNs at 48 hours post-fertilization (hpf) in embryos embedded in low-melt agarose (**Fig. 4A**). To determine whether *C9orf72* KO affected PMNs organization, we first measured the distance between the common PMNs ventral root and the longest ventral axon projection of the first five hemisegments caudal to the yolk (**Fig. 4A, B**). The distance between PMNs ventral roots was unaffected by *C9orf72* KO at 48 hpf, as was the longest ventral projection length (**Fig. 4D, E**). We next investigated whether *C9orf72* KO induced more subtle morphological defects in PMNs of larvae. To do so, we manually traced the axonal filaments of PMNs in 3D within the first two hemisegments caudal to the yolk (**Fig. 4A, C**), starting at the common axonal ventral root of PMNs. Our analysis revealed that *Hb9^-/+^C9orf72^-/-^* larvae exhibited mild axonal disruption in PMNs, characterized by a slight but statistically significant increase in axonal branching at 48 hpf compared to *Hb9^-/+^C9orf72^+/+^* controls (*p = 0.0332*; 46.48 ± 0.70 vs. 43.86 ± 0.95 branches per millimeter of filament) (**Fig. 4F**). Additionally, *C9orf72* KO specimens displayed a significantly greater total axonal filament length in the region of interest (*p < 0.001*; 3497 ± 119 µm vs. 2970 ± 60 µm).

**Fig 4.**
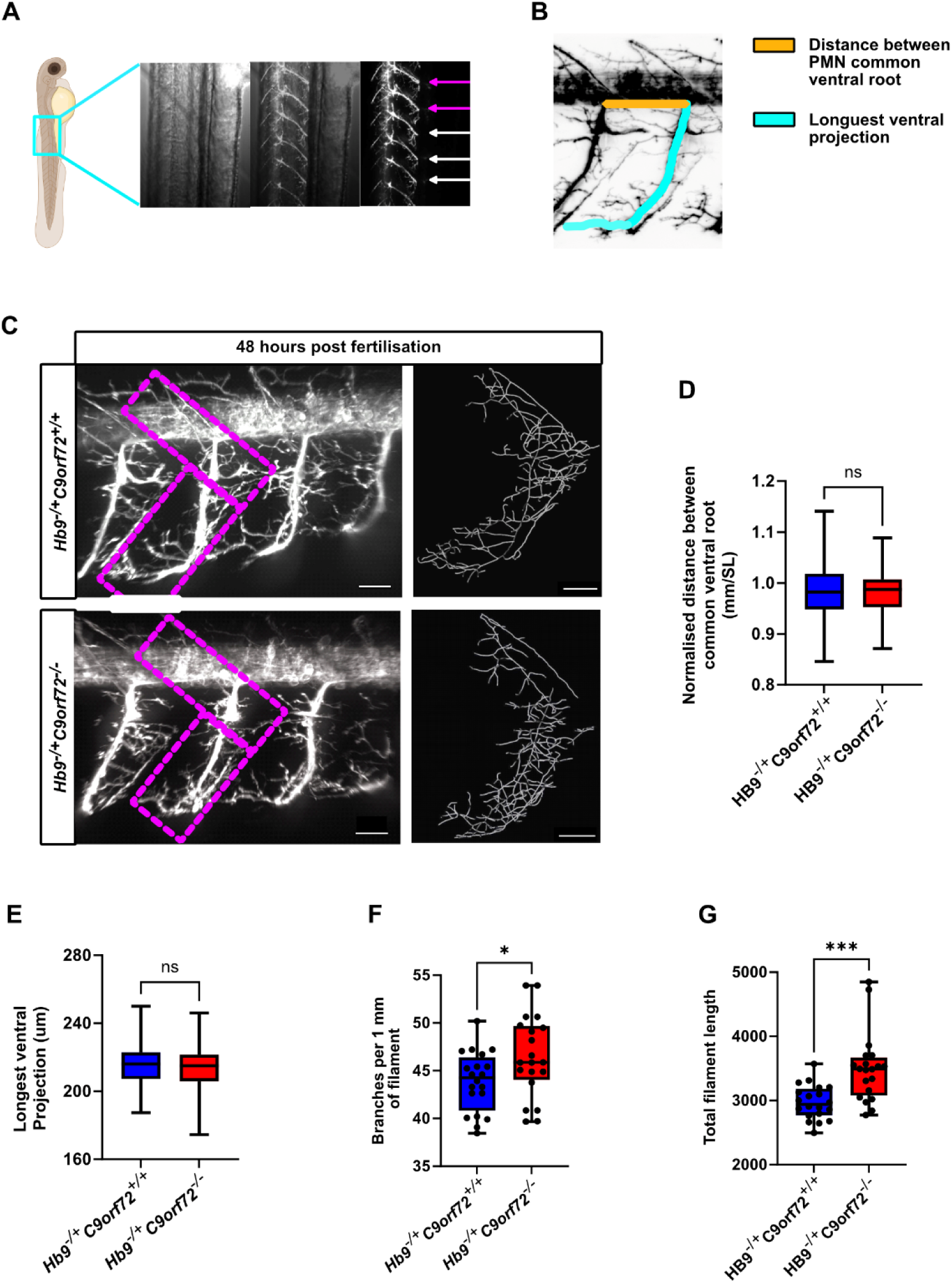
Complete loss of C9orf72 leads to mild axonal over-branching of spinal primary motor neurons. **(A)** Schematic representation of the dorsal primary motor neuron (PMN) units analyzed for measurements of the common ventral root distance (VRD), longest ventral axonal filament (LVF), and 3D axonal filament tracing in *Hb9^-/+^C9orf72^+/+^* and *Hb9^-/+^C9orf72^-/-^* larvae at 48 hours post-fertilization (hpf). **(B)** Schematic depiction of VRD and LVF measurements, illustrating how the distance between the common PMN’s ventral root and the longest projecting axonal filament was quantified. **(C)** Representative confocal image of a spinal hemisegment PMN unit and corresponding 3D axonal filament tracing used to assess axonal branching. Dashed lines indicate the area considered for filament tracing. **(D, E)** Quantitative analysis of VRD **(D)** and LVF **(E)** revealed no significant differences between *Hb9^-/+^C9orf72^+/+^* and *Hb9^-/+^C9orf72^-/-^* larvae. To account for size differences, VRD was normalized to the mean hemisegment length of the five considered for each specimen (N = 20, n = 100). **(F, G)** Quantitative analysis of axonal branching showed a mild but significant increase in the number of branches per 1 mm of filament **(F)** and a significant increase in total filament length **(G)** in *Hb9^-/+^C9orf72^-/-^* larvae compared to controls (N = 10, n = 20). *Statistical tests:* Unpaired two-tailed parametric t-test **(D–F**); Mann-Whitney test **(G)**. **\***\* * *p ≤ 0.001*, * *p ≤ 0.05, NS p > 0.05.* Boxplot representation: Extremities indicate minimum and maximum values; box limits represent the interquartile range (central 50%), and the central line marks the median. N represents the number of distinct specimens; n represents the total number of PMN units per genotype analyzed. Scale bars = 30 µm.

To further investigate potential neurological defects underlying the motor deficits observed in *C9orf72* KO larvae, we examined NMJ integrity in spinal hemisegments. We performed double immunohistochemistry on 6 dpf larvae, using an SV2 antibody to mark presynaptic terminals and α-bungarotoxin (α-BTX) to label postsynaptic acetylcholine receptors. Quantitative analysis revealed a significant increase (Ordinary one-way ANOVA, *p < 0.001*) in the total number of postsynaptic α-BTX puncta in *C9orf72^-/-^* larvae (4752 ± 133.4 puncta) compared to *C9orf72^+/+^* (3958 ± 164.5 puncta) and *C9orf72^-/+^* specimens (3842 ± 149.3 puncta) (**S6 Fig. B**). However, there was no significant difference in the total number of presynaptic SV2 puncta (**S6 Fig. 4C**) or in the number of colocalized presynaptic and postsynaptic puncta at either the 100% or 50% colocalization threshold (**S4 Fig. D-G**). These findings suggest that complete loss of C9orf72 selectively alters the postsynaptic compartment at the NMJ, while overall synaptic architecture and alignment remain unaffected.

### Therapeutic compound screening and validation

The motor deficits observed in our *C9orf72* KO zebrafish model represent an ALS-related phenotype suitable for therapeutic compound screening. However, zebrafish are less amenable to high-throughput screening than *C. elegans*. Therefore, large compound libraries can first be screened in *C. elegans* utilizing ALS-related phenotypes, with candidates refined through validation of additional phenotypes and the most promising hits tested in our zebrafish model. Notably, *C. elegans* display a stereotyped swimming motion in liquid medium that actively engages the NMJ, making it an effective model for assessing MN health [33]. Additionally, high-throughput liquid culture assays using automated infrared beam scattering have been developed to measure locomotor activity and evaluate the neuroprotective effects of compounds in *C. elegans*. Given that *alpha-1(ok3062)* mutants exhibit MN degeneration and NMJ dysfunction, we investigated their locomotor phenotype in liquid culture to confirm their suitability for high-throughput screening. After 30 minutes, young adult *alpha-1(ok3062)* mutants showed significantly reduced swimming activity compared to wild-type (N2) worms (*p = 0.0015*, **Fig. 1A**). This deficit persisted for up to six hours (**Fig. 5A**) and was suitable for high-throughput compound screening. We screened over 4,000 bioactive compounds from commercially available drug libraries using a high-throughput liquid assay, identifying 80 compounds that significantly enhanced locomotion in *alpha-1(ok3062)* mutants (*p < 0.05*, **S3 Table**). Of these, the top 12 compounds were selected for further validation on solid media, all of which significantly improved age-dependent paralysis and neurodegeneration phenotypes in *alpha-1(ok3062)* mutants (*p < 0.05*, **S4 Table**) [34].

**Fig 5.**
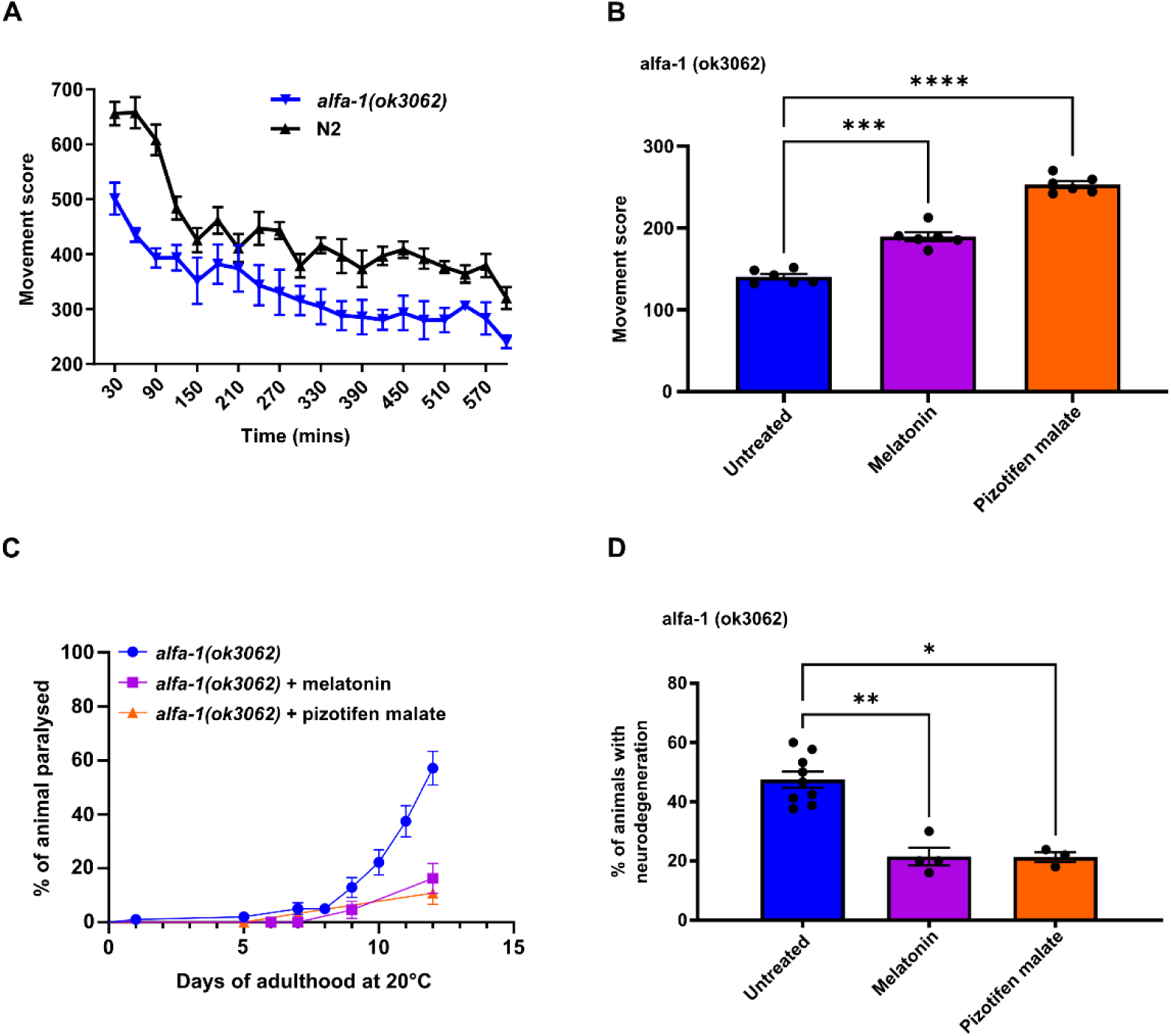
Pizotifen malate and melatonin ameliorate deleterious phenotypes in *C. elegans knockout for C9orf72* orthologue (*alfa-1*). **(A-B)** Motor activity of *alfa-1(ok3062)* and N2 (wild-type) worms measured in liquid medium using automated infrared beam scattering. (A) *alfa-1(ok3062)* worms exhibit motility impairment after 30 minutes compared to N2 controls, which persists throughout the 10-hour assay. Data are presented as mean ± SEM; n = 3 technical replicates. **(B)** Improvement in motor activity of *alfa-1(ok3062)* worms treated with melatonin or pizotifen malate (PM) compared to untreated controls. Motor activity was measured in 30-minute bins over 180 minutes. Data are presented as mean ± SEM; N = 3 technical replicates, n = 6 repeated measurements per replicate. Two-way ANOVA with Dunnett’s post-hoc test. *** *p < 0.001*, **** *p < 0.0001*. **(C)** Kaplan-Meier survival analysis of age-dependent paralysis in *alfa-1(ok3062)* worms on solid media containing 1% DMSO (untreated), melatonin, or PM. Both compounds significantly delay paralysis compared to untreated controls. *p < 0.001* (log-rank Mantel-Cox test); N = 3 plates, n = 60-99 worms per condition. **(D)** Percentage of *alfa-1(ok3062)* worms displaying neurodegeneration at 9 days post-adulthood on solid media containing 1% DMSO (untreated), melatonin, or PM. Both compounds significantly reduce neurodegeneration. Kruskal-Wallis test with Dunn’s multiple comparison post-hoc test. * *p < 0.05*, ** *p < 0.01*; N = 3-9 worm synchronisation, n = 90-270 worms per condition. Data are presented as mean ± SEM.

Among the 12 compounds, melatonin and pizotifen malate (PM) stood out for their effectiveness on *alfa-1(ok3062)* worm ALS-related phenotypes, as well as their accessibility and previous use in zebrafish larvae [35,36]. Both compounds improved locomotor deficits, with treated mutant worms showing significantly greater locomotor activity (melatonin: 189.4 ± 5.3 units; PM: 253.1 ± 4.3 units) compared to untreated controls (140.4 ± 3.4 units; *p < 0.001* for both, **Fig. 5B**). They also increased the proportion of *alfa-1(ok3062)* mutants unaffected by age-related paralysis at day 12 (melatonin: 83.7 ± 5.6%; PM: 89.1 ± 4.3%) compared to untreated controls (42.9 ± 6.3%; *p < 0.001* for both, **Fig. 5C**). Neurodegeneration at day 9 was also significantly reduced in treated *alfa-1(ok3062)* worms (melatonin: 21.3 ± 3.0%; PM: 21.27 ± 1.3%) compared to untreated controls (47.5 ± 2.7%; *p = 0.0078* and *p = 0.0313*, respectively, **Fig. 5D**). Therefore, we aimed to validate the therapeutic potential of melatonin and PM for C9orf72 LOF-related phenotypes in zebrafish. To do this, we assessed their ability to alleviate swimming deficits seen in *C9orf72* LOF larvae.

We exposed 6 dpf *C9orf72* KO zebrafish larvae to melatonin or PM through in-water dosing that started 2 hours post-fertilization, and we measured their swimming activity under a 120-minute light-dark paradigm (**Fig. 6A**). While various concentrations of melatonin did not significantly improve motor activity in any of the *C9orf72* LOF genotypes (*p > 0.05*, **S7 Fig.**), PM treatment at a relatively low concentration significantly enhanced total swimming activity in *C9orf72^-/-^* larvae (7003 ± 255 mm vs. 10,380 ± 438 mm; p < 0.0001, **Fig. 6B, E**). PM also increased total swimming activity in wild-type (*C9orf72^+/+^*) larvae (11,592 ± 379 mm vs. 13,628 ± 541 mm; p = 0.0075, **Fig. 6B, C**). In contrast, the increase in total swimming activity for *C9orf72^-/+^* larvae due to PM exposure was not statistically significant (**Fig. 6B, D)**. The extent of PM-induced locomotion increase was greater *in C9orf72^-/-^* larvae (+38.9 ± 0.06%) than in wild-type larvae (+16.2 ± 0.05%). The effects of PM were especially pronounced during the dark phases, with generally weaker effects during the light phases (**Fig. 6C-E**).

**Fig. 6.**
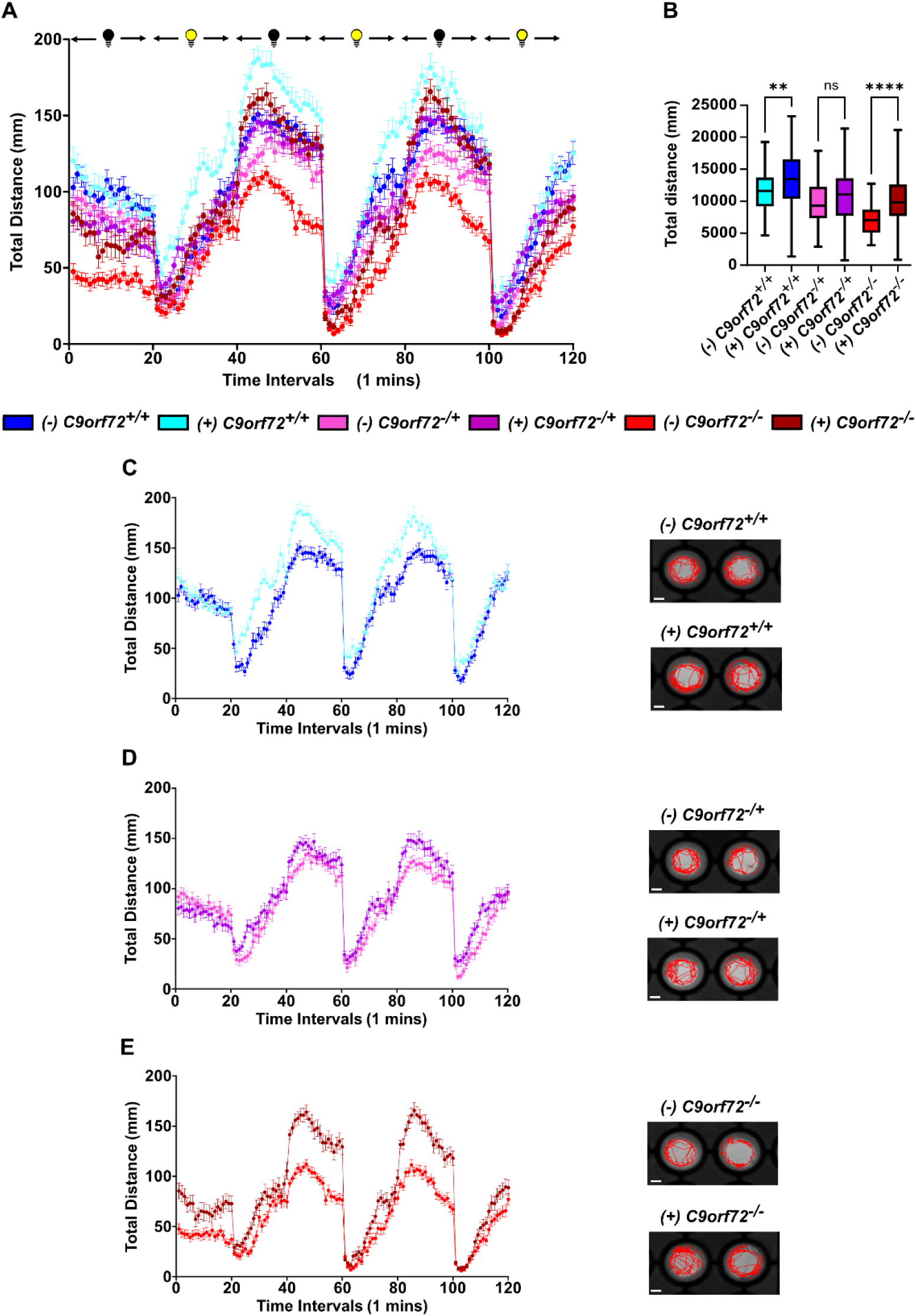
Pizotifen malate treatment increases total swimming activity in controls and partially rescues swimming deficits in C9orf72 LOF zebrafish larvae. **(A)** Mean swimming distance per minute for pizotifen malate (PM) treated (+) and untreated (-) *C9orf72^+/+^*, *C9orf72^-/+^*, and *C9orf72^-/-^ six*-day post-fertilization (dpf) zebrafish larvae under a 120-minute phasic dark-light program. Data are presented as mean ± SEM. **(B)** Quantitative analysis of total swimming distance shows significant increases in PM-treated (+) *C9orf72^+/+^* and *C9orf72^-/-^* larvae compared to untreated (-) controls. Data are presented as mean ± SEM. Welch and Brown-Forsythe ANOVA with Dunnett’s T3 multiple comparisons post-hoc test. **** *p < 0.0001*, ** *p ≤ 0.01*, NS *p > 0.05*. Boxplot extremities indicate minimum and maximum values; box limits represent the central 50% of the data, and the central line marks the median. **(C-E)** Swimming distance per minute and representative 60-second swimming tracks for treated and untreated *C9orf72^+/+^* (C), *C9orf72^-/+^* (D), and *C9orf72^-/-^* (E) larvae. Data are presented as mean ± SEM. N = 5, experimental repeats; n = 80, larvae per genotype. Scale bars = 0.2 cm.

## Discussion

### Insights from a new C9orf72 loss-of-function zebrafish model

*C9orf72* hexanucleotide repeat expansions are the most common genetic cause of ALS/FTD. It remains unclear whether *C9orf72* loss-of-function, DPR production, and RNA foci gain-of-function, or a combination of both, is causative for the disease. Here, we generated and characterized a new stable *C9orf72* KO zebrafish line, providing a valuable vertebrate model to investigate *C9orf72* LOF and validate potential therapeutic compounds through its *C9orf72*-related ALS phenotypes. The *C9orf72^-+/+^* and *C9orf72^-/-^* mutants exhibited motor deficits, growth delays, and increased mortality from early to late larval stages. *C9orf72^-/-^* larvae also showed mild overbranching and increased axonal filament length in spinal PMNs, along with subtle NMJ disruptions characterized by increased postsynaptic acetylcholine receptor density. Despite these deficits, *C9orf72^-/-^* specimens developed normally into adulthood, surviving and reproducing for up to 12 months without significant health deterioration. These findings support a role for C9orf72 deficiency in the manifestation of *C9-*ALS related phenotypes but suggest that LOF alone may be insufficient to induce overt neurodegenerative phenotypes in zebrafish during early larval stages.

### Comparison to previous C9orf72 loss-of-function zebrafish models

RNAi-based KD of *C9orf72* in zebrafish larvae has been shown to cause severe overbranching and truncation of caudal primary MNs [17], as well as motor deficits, morphological abnormalities, reduced survival, and disrupted NMJ integrity [18]. In contrast, our *C9orf72^-/+^* larvae displayed no overt morphological defects, only a mild reduction in survival, and a persistent but less pronounced decrease in motor activity. The *C9orf72^-/-^* specimens exhibited early mild overbranching of spinal PMN axons, with longer axonal filaments primarily due to increased branching and a greater number of postsynaptic structures. While *C9orf72^-/-^* specimens exhibited more pronounced survival and motor deficits than *C9orf72^-/+^* specimens, these phenotypes were still milder than those reported in the microRNA-based *C9orf72 KD* model. Unlike RNAi-based approaches, our zebrafish *C9orf72* KO model incorporates a stable genomic mutation, reducing susceptibility to off-target effects but potentially triggering compensatory genetic responses via nonsense-mediated decay [37]. Additionally, it enables the investigation of complete C9orf72 loss effects. The phenotypic discrepancies between our *C9orf72^-/+^* specimens and the RNAi KD models highlight the challenges in interpreting KD-based findings without validation in a stable LOF mutant.

The recently reported CRISPR/Cas9-generated *C9orf72* KO zebrafish model supports several of our findings. Notably, they observed an absence of both widespread NMJ alterations and MN degeneration in the spinal cord of their adult specimens [19]. Our C9orf72 KO model does not phenotypically conflict with the previously reported KO model, although there is limited overlap in the scope and nature of the characterization approaches. A key distinction between the two KO models is the predicted truncated C9orf72 protein, which is 29% in their model compared to 15% in ours, resulting from a premature stop codon downstream of our mutation sites. Furthermore, we confirmed complete C9orf72 protein ablation in *C9orf72^-/-^* specimens using a known cross-reactive antibody for human and mouse, which we validated for zebrafish C9orf72, whereas their model demonstrated reduced *C9orf72* mRNA levels [19].

### Visual Impairment as a possible confounding factor

Jaroszynska et al. (2024) reported retinal degeneration and gliosis in 24-month-old *C9orf72^-/-^* zebrafish, indicating potential visual impairment. However, we argue that this effect is minimal during larval stages, as they found no retinal abnormalities at 5 days post-fertilization (dpf), and only limited inner nuclear layer neuron loss was identified by 8 months, with the photoreceptors remaining unaffected. Additionally, locomotor deficits in *C9orf72^-/-^* and *C9orf72^-/+^* larvae continued even under conditions that reduce the impact of acute light stimuli, such as our biphasic dark-light paradigm and locomotor assessment conducted only after light adaptation (SSWAP). Furthermore, a zebrafish model of moderate perceptual alteration (hyperopia) showed no changes in spontaneous swimming distance at 7 dpf under consistent light or dark conditions [38]. Altogether, these findings suggest that neuromuscular disruptions or other *C9orf72* loss-of-function effects, rather than visual impairment, primarily drive the locomotor phenotype in our zebrafish model.

### Pizotifen malate as a potential therapeutic compound for ALS

We demonstrated that *alfa-1(ok3062) C. elegans* exhibit early motor deficits in liquid, which are suitable for automated high-throughput compound screening. This enabled us to identify 80 compounds that improved the locomotor deficits, with 12 also demonstrating the ability to alleviate age-dependent paralysis and neurodegeneration. Among the 12, melatonin and PM were selected for further phenotypic compound validation in *C9orf72* loss-of-function zebrafish larvae by evaluating their persistent locomotor deficiencies. Locomotion assays revealed that low-dose PM (0.5 µM) significantly improved swimming deficits in *C9orf72^-/-^* larvae at 6 days post-fertilization, while melatonin did not show significant effects across genotypes or tested concentrations.

PM is approved for human use in Canada [39], France [40], and the UK [41]. It primarily acts as a 5-HT₂ serotonin receptor antagonist with weak anticholinergic, H1 antihistamine, and antikinin activity. Historically, PM has been utilized for preventing vascular headaches [42] and treating depressive disorders [43]. Given the existence of zebrafish orthologues for 5-HT₂ [44] and H1 receptors [45], higher PM doses were anticipated to induce mild sedation, consistent with findings in rodents and primates [42]. At 5 µM, PM decreased locomotion in *C9orf72^+/+^* and *C9orf72^+/-^* larvae but did not affect locomotion in *C9orf72^-/-^* larvae, whereas 0.5 µM surprisingly increased locomotion in *C9orf72^+/+^* larvae. This effect may indicate adaptive changes in neurotransmitter release, receptor sensitivity, or compensatory excitatory mechanisms, potentially leading to a rebound increase in locomotion as PM’s inhibitory effects fade over time. PM’s capacity to alleviate paralysis and neurodegeneration in *C. elegans alfa-1 (ok3062)* mutants implies that its effects in *C9orf72* KO zebrafish involve both neuroprotective and compensatory mechanisms. This interpretation is supported by the relatively larger increase in locomotion in *C9orf72^-/-^* larvae (1.482-fold) compared to *C9orf72^+/+^* larvae (1.176-fold). These findings are consistent with previous reports of PM reducing neurodegeneration and enhancing motor performance in the R6/2 Huntington’s disease mouse model [46].

## Conclusion

We generated and characterized stable *C9orf72^-/+^* and *C9orf72^-/-^* zebrafish lines, whose phenotypes support a link between *C9orf72 loss of* function (LOF) and the manifestation of ALS-related phenotypes. However, they demonstrated significant discrepancies with the known zebrafish RNAi-based *C9orf72* knockdown (KD) models. These lines serve as valuable tools for studying C9orf72 function, the synergistic effects of ALS risk factors, mutations, and pathways potentially causative for the pathology, as well as for validating prospective ALS therapeutic compounds in a rapid and cost-effective *in vivo* vertebrate model system. We showed that the *C. elegans* alfa*-1(ok3062)* mutants exhibit a locomotor deficit phenotype suitable for high-throughput compound screening, resulting in the identification of 12 compounds that demonstrated neuroprotective effects and improved locomotion. Finally, PM’s ability to alleviate *C9orf72* deficiency-related ALS locomotion phenotypes was validated in the *C9orf72^-/-^* specimens, suggesting its potential for further validation in other ALS models as a prospective therapeutic for the disease. Further validation of PM’s potential to alleviate ALS-related phenotypes, particularly those involving neurodegeneration, in other vertebrate ALS models will be essential for clinical translation.

## Material and methods

### Fish husbandry

All *Danio rerio* lines were maintained at 28.5°C under a 12-hour light/dark cycle and staged according to Swaminathan et al. (2018). All experiments were conducted at the Centre de Recherche du Centre Hospitalier de l’Université de Montréal (CRCHUM) in compliance with the Canadian Council for Animal Care (CCAC) guidelines and approved by the Comité Institutionnel de Protection des Animaux (protocol #N21017APz). Most experiments were performed on sexually undifferentiated larvae between 2 and 20 dpf, except for extractions conducted on 6-month-old female specimens. Humane endpoints were established, and animals were monitored daily for well-being, as per guidelines established by the Canadian Council of Animal Care committee at CRCHUM. Adult zebrafish were euthanized if they exhibited severe health deterioration, including an inability to feed or swim. Physical abnormalities such as distended abdomen, skin ulcerations, skeletal deformities, or severe swimming impairments were also monitored daily. Affected adults were euthanized via rapid chilling (hypothermal shock) immediately upon detection of severe health decline.

### sgRNA and Cas9 preparation for C9orf72 knockout line generation

The single-guide RNA (sgRNA) sequence was designed using CRISPRscan to target an early coding region in exon 2 of the C9orf72 gene (ENSDARE00000573949), with the protospacer adjacent motif (PAM) sequence noted in parentheses: GCGCAGCGCAGAGAGC<u>GG</u>CG (CGG). The synthesis of sgRNA and Cas9 mRNA followed the protocols described by Moreno-Mateos et al. (2015). Microinjections were conducted in Tübingen Long Fin (TL) wild-type zebrafish embryos in accordance with Samarut et al. (2016). The most likely off-target sites were predicted using CRISPRoff v1.1 [28].

### Genotyping

#### Primer design and selection

HRM, PCR and sequencing primers were designed with SnapGene (Dotmatics) version 4.2.1.1 in conjunction with the online tool Primer-BLAST [50] The second exon of *C9orf72* was sequenced to characterize indels using the following primers: F 5’ GCCAAGACGAAG AACTTGACATCC, R 5’ GGAACAATCTCGGATGACAAC. and following the general guidelines established by Chuang, L. Y. et al (2013). All primer sets are available upon request.

#### Fin clip sample collection and DNA extraction

Adult zebrafish were anesthetized in tricaine methanesulfonate (MS222; Sigma-Aldrich) at 160 mg/L, and a small caudal fin sample was excised using a sharp blade. Fish were immediately transferred back to fresh water in isolated tanks to recover. Genomic DNA extraction was performed in 20 μL of 50 mM NaOH, followed by boiling for 10 minutes. The reaction was then buffered by adding 1/10 volume of 100 mM Tris-HCl (pH 8.0).

#### High-resolution melting (HRM)

Identification of the presence of indels in the 2^nd^ exon of *C9rof72* following the injections and genotyping of the specimen from the *C9orf72* KO line was done using the F 5’ CGGAGAGGTCACATTTCTGGCC and R 5’ GCCAAGACGAAGAACTTGACATCC primers. Integration of indels was identifiable by shift in the Δ Fluorescence/ Δ temperature HRM curve profile and the genotyping was done by matching curve profiles of tested specimens to those of specimens whose genotype was confirmed by sequencing.

The PCR reactions were performed as described by Samarut et al. (2016) in a LightCycler 96 (Roche). HRM curves were analyzed using the Roche LightCycler 96 software (version 1.1).

#### PCR and sequencing

Second exon sequencing of *C9orf72* for characterization of the indels were done using: F 5’ GCCAAGACG AAGAACTTGACATCC and R: 5’ GGAACAATCTCGGATGACAAC. Sequencing of the 3^rd^ intron of pou3f2b to validate the absence of off-target indels was done using F 5’ GTCAAATCACCAAACCACC ACCCA and R 5’ AGGTGTTCGCAGAAAGCATTGC. Sequencing of the 2^nd^ intron of *SI:DKEY-277I15.2* to validate the absence of off-target indels was done using F 5’ GCACATGTAGACACTTCGCCTCT and R 5’ ACACATGCTCAATCTTCGTTCTCC. Sequencing of the 11^th^ exon of *ch25hl3* to validate the absence of off-target indels was done using F 5’ GTGCATTGCTTGCTGATGCTAAG, R 5’ CACCGACTGGCAC AATGTAGTC.

The PCR reactions were made with 0.5 μL of dNTP (10 μM), 0.5 μL of each primer (10 μM), 2.5 μL of 10x PCR buffer, 0.125 μL of Taq DNA polymerase (GenedireX), 1 μL of genomic DNA and water up to 25 μL. The PCR reaction protocol was 94 °C for 5 min, then 35 cycles of 94 °C for 30 s, 57-60 °C for 30 s and 72 °C for 45 s and finally 72 °C for 10 min. Samples were sequenced by the Genome Quebec/McGill center using Applied Biosystems 3730xl DNA Analyzer.

### C9orf72 knockout validation

#### TaqMan gene expression analysis

To assess *C9orf72* relative transcript expression, we used TaqMan Gene Expression Assays (Applied Biosystems, Thermo Fisher Scientific) with the following probes:

zgc:100846 (C9orf72): Dr03094731_m1

Polr2d (housekeeping gene): Dr03095551_m1

Total RNA was extracted from 30 pooled larvae (6 dpf) using TriReagent® (Sigma-Aldrich) according to the manufacturer’s protocol. RNA was quantified using a NanoPhotometer (Implen) and stored at −80 °C until further use. cDNA was synthesized from 1 µg of total RNA using the Superscript VILO cDNA Synthesis Kit (Thermo Fisher Scientific). Undiluted cDNA was used for real-time PCR, yielding CT values between 25 and 32. Gene expression was quantified using a QuantStudio 3 Real-Time PCR System (Thermo Fisher Scientific) and analyzed using the ΔΔCT method, with Polr2d as the normalization control.

#### Western blot

Protein samples for SDS-PAGE and Western blot (WB) analysis were prepared as follows:

Embryo Samples: At 48 hours post-fertilization (hpf), 40 zebrafish embryos per condition were manually dechorionated. Embryos were lysed and homogenized using a pellet pestle in 150 µL of ice-cold lysis buffer containing 150 mM NaCl, 1.0% Triton X-100, 0.1% SDS, 50 mM Tris (pH 7.5), 0.5 mM EDTA, and a protease inhibitor cocktail (1:10, Sigma-Aldrich). Lysates were then boiled for 5 min, centrifuged at 16,000 × g for 10 min at 4°C, and the supernatant was collected. Protein concentration was measured using the Bradford assay (BioRad), and samples were stored at −70°C until further analysis.

Adult Brain Samples: Adult zebrafish (6 months old) were euthanized by prolonged immersion in 300 mg/L MS-222 (Tricaine mesylate; Sigma-Aldrich). Specimens were rinsed with phosphate-buffered saline (PBS) and decapitated using a razor blade. Brains were surgically dissected, pooled (4 brains per condition) into separate 2 mL Eppendorf tubes, weighed, and homogenized on ice with a pellet pestle in 60 µL of ice-cold lysis buffer per 100 mg tissue until no visible tissue remained. Further homogenization was performed by sonication on ice (6 cycles, 20% amplitude, 5 seconds on, 10 seconds off). Subsequent processing steps were identical to those used for embryo samples.

Western Blotting: For Western blot analysis, 85 µg of protein lysate per sample was loaded onto 5–16% gradient polyacrylamide gels and transferred to nitrocellulose membranes. Ponceau S staining was used to visualize transferred proteins. Membranes were blocked in 5% milk in TBST (TBS with 0.1% Tween-20) and incubated overnight at 4°C with the C9orf72 primary antibody (ab221137, 1:2500 dilution in 5% BSA/TBST). The following day, membranes were incubated for 1 hour at room temperature with peroxidase-conjugated secondary antibodies (1:10,000 dilution in 5% milk/TBST), followed by multiple washes. Detection of immunoreactive bands was performed using a LI-COR Odyssey Imaging System (LI-COR Biosciences), and data analysis was conducted with LI-COR Image Studio Lite Version 5.2.

### C9orf72 knockout characterization

#### Morphological measurement and gross morphology assessment

Standard length and morphological phenotypes were assessed in larvae at 2, 4, 6, 10, 15, and 20 dpf. For each time point, larvae from three independent batches (each containing 20–30 larvae per genotype) were analyzed. For imaging, larvae were individually anesthetized in tricaine methanesulfonate (MS222; Sigma-Aldrich) at a final concentration of 160 mg/L in E3 medium (5 mM NaCl, 0.17 mM KCl, 0.33 mM CaCl₂, 0.33 mM MgSO₄, 0.00001% w/v Methylene Blue, pH 7.2). Larvae were then mounted in 0.5% (w/v) low-melting-point agarose (Ultrapure™ LMP Agarose; Invitrogen) in E3 medium on 6 cm petri dishes. Before imaging, dishes were filled with E3 medium to fully submerge the mounted larvae. Lateral images were acquired using a Leica S6E stereomicroscope equipped with an iPhone 6 mounted on an iDu Optics cast from LabCam. Standard body length was measured from the anterior tip to the caudal peduncle using ImageJ software.

#### Survival assay

At 24 hpf, 100 embryos per genotype with normal growth and no visible defects were transferred to 500 mL tanks containing 250 mL of E3 medium and maintained in a dry incubator, as described in the fish husbandry section. Two-thirds of the E3 medium was replaced daily with fresh medium. Larvae were fed twice daily, and dead specimens were recorded and removed before each feeding. Every two days, larvae were transferred to clean 500 mL tanks to maintain optimal water quality.

#### Swimming activity monitoring assay

At 6, 8, 10, 12, 15, and 20 dpf, larvae (24 per group for *C9orf72^+/+^* and *C9orf72^-/-^*, 48 for *C9orf72^-/+^* mutants) were individually transferred to a 96-well plate containing fresh E3 medium. Swimming activity was recorded under two different paradigms. Phasic paradigm: recording was conducted over a 120-minute alternating dark-light cycle, consisting of six 20-minute phases. Biphasic paradigm: recording was done over an 80-minute alternating dark-light cycle, comprising a 20-minute dark phase followed by a 60-minute light phase. For both paradigms, larvae were placed in a dark recording chamber for 20 minutes to acclimate before recording began. Recordings were performed using a Basler GenIcam camera within a DanioVision recording chamber (Noldus). Ethovision XT 12 (Noldus) was used to quantify total swimming distance, with mean swimming distance per minute and cumulative swimming distance for the entire assay exported for further analysis in GraphPad Prism. Swimming distance was normalized to the mean BL measurement at the corresponding time point for each genotype.

#### Spinal primary motor motoneurons morphology evaluation assay

Transgenic *Hb9:GFP* zebrafish (*Tg(mnx1:GFP)*) were crossed with *C9orf72^-/-^* zebrafish to obtain *Hb9:GFP C9orf72^-/+^* specimens. A stable *Hb9:GFP C9orf72^-/-^* line was established through multiple incrosses. For controls, transgenic *Hb9:GFP* zebrafish were crossed with wild-type TL specimens to generate *Hb9:GFP C9orf72^+/+^*, and *Hb9:GFP C9orf72^-/+^* were crossed with *C9orf72^-/-^* to ensure that all GFP-positive embryos were heterozygous for *Hb9:GFP*.

At 30 hours post-fertilization (hpf), embryos expressing GFP in the spinal cord were selected. At 48 hpf, selected embryos were anesthetized using tricaine methanesulfonate (MS222, Sigma-Aldrich) at a final concentration of 160 mg/L in E3 medium. Once immobilized, embryos were embedded in 0.5% (w/v) low-melting-point agarose (*Ultrapure™ LMP Agarose, Invitrogen*). To visualize spinal primary motor neurons (PMNs) axonal architecture, embryos were imaged using confocal microscopy on an Olympus BX61W1 microscope equipped with a Quorum Technology (Ontario) spinning disk head and a Hamamatsu ORCA-ER camera. The first two PMNs units caudal to the yolk were imaged to produce z-stacks (85–90 slices) at 20× magnification, with an x/y resolution of 512 × 512 pixels (pixel size: 0.33 µm) and a z-resolution of 1 µm. Images were acquired using Volocity (version 7.0, Improvision) and analyzed using Imaris (version 8.1.1) with its filament tracing application.

#### Neuromuscular junction integrity evaluation assay

At 6 days post-fertilization (dpf), 20 zebrafish larvae per genotype were fixed overnight at 4°C in 4% paraformaldehyde (PFA) in 1× PBS. The next day, samples were washed three times (3×) for 15 min in 0.1% Tween-20 in PBS (PBS-Tween) at room temperature (RT). To permeabilize tissues, larvae were incubated in 1 mL of 1 mg/mL collagenase (Sigma-Aldrich, C0130-100MG) in PBS for 150 min at RT on a rocker. Afterward, samples were rinsed 3× in 1% Triton X-100 in PBS (PBS-Triton) for 10 min at RT on a rocker. For blocking, larvae were incubated in blocking solution (1% bovine serum albumin, 1% DMSO, 1% Triton X-100, 2% normal goat serum in PBS) for 1 h at RT. To label postsynaptic acetylcholine receptors, larvae were incubated in 10 mg/mL tetramethylrhodamine-conjugated α-bungarotoxin (ThermoFisher, T1175) in 0.1% PBS-Tween for 30 min at RT. After three 15-min washes in PBS-Tween, larvae were incubated overnight at 4°C in blocking solution containing the primary antibody SV2 (1:200, Developmental Studies Hybridoma Bank) to stain presynaptic vesicles. The next day, samples were washed 3× in PBS-Tween for 15 min and incubated overnight at 4°C in 1:1000 Alexa Fluor 488 goat anti-mouse antibody (Jackson ImmunoResearch, 115-545-205) in blocking solution. After a final 3× PBS-Tween wash (15 min at RT), larvae were stored in 80% glycerol in PBS at 4°C after 1 h at RT on a shaker. For microscopy preparation, larvae were surgically decapitated and mounted in 80% glycerol on 25 × 75 × 1.0 mm microscope slides under a 24 × 50 mm coverslip, with petroleum jelly grease bridges to prevent compression.

The integrity of neuromuscular junctions (NMJs) in spinal motoneurons was analyzed using confocal microscopy on a Zeiss spinning disk Axio Observer Z1 (Carl Zeiss, Germany). For each larva, Z-stacks (100–115 slices) from the first three spinal hemisegments caudal to the yolk were obtained at 40× magnification with the following settings: x/y resolution: 512 × 512 pixels (0.33 µm per pixel); z-resolution: 0.44 µm; Tiling: Applied (x or y axis = 2) to capture the full hemisegment; stitched images were employed for analysis. Images were captured using ZEN 2.6.7 Blue edition (Improvision, England). NMJ integrity analysis was conducted using FIJI/ImageJ (version 2.9.0/1.53t) with the “NMJ Analyser” macro (Version 8.11) under default settings, as outlined in Singh et al. (2023).

### Drug validation assay

For all conditions, chemical treatments were administered through in-water dosing, starting at the 64-cell stage (2 hours post-fertilization, hpf) and continued until 6 days post-fertilization (dpf). Vehicle-treated embryos (0.01% dimethyl sulfoxide [DMSO]) served as controls. The compounds tested were pizotifen malate (PM, Cayman Chemical; #20765-500) and melatonin (Cayman Chemical; #14427). Stock solutions were made by dissolving PM (30 mM) and melatonin (10 mM) in 100% DMSO. Working solutions for PM, melatonin, and DMSO vehicle (0.01%) were prepared through sequential dilution in E3 medium and added directly to the wells of a 12-well plate containing 20 zebrafish embryos per well. The medium was refreshed daily to ensure compound stability and minimize degradation. To validate drug effects, swimming activity assays were conducted as previously described. Before the assay, all medium was replaced with fresh E3 medium without DMSO, PM, or melatonin to eliminate any residual effects of acute exposure.

### C. elegans

#### Strains and maintenance

Standard methods for culturing and handling the worms were employed [53]. Worms were cultured on standard nematode growth medium (NGM) streaked with OP50 *Escherichia coli*. The stock population was maintained at 15 °C, while experimental worms were kept at 20 °C unless specified otherwise. N2 (wild-type), RB2260 (*alfa-1(ok3062) II)*, EG1285 (*oxIs12 [unc-47p::GFP + lin-15(+)])* and a strain resulting from the cross of RB2260 and EG1285 (*alfa-1(ok3062) II; lin-15B&lin-15A(n765) oxIs12 [unc-47p::GFP + lin-15(+)] X*) were utilized. All experiments were conducted on hermaphrodites. Most strains were supplied by the CGC, which is supported by the NIH Office of Research Infrastructure Programs (P40 OD010440).

#### Paralysis assay

Day one adult worms were transferred on 5μM fluorodeoxyuridine (FUDR; Sigma-Aldrich) plates. Worms were scored daily for movement for 12 days. Worms were counted as paralyzed if they failed to move after prodding on the nose. Experiments were performed at 20 °C and at least 60 worms were counted per condition.

#### Liquid culture assay

A synchronized population was obtained using hypochlorite extraction. Worms were grown on solid media until day 1 of adulthood. On day 1, 50 worms per well were placed in S basal with OP50 *E. coli* (optical density 0.5) in a flat-bottom 96-well plate. Standard errors are displayed on the graph. Measurements were taken using a Microtracker (Phylumtech) with standard parameters for *C. elegans*.

#### Neurodegeneration

Animals were selected for in vivo visualisation on days 1, 5, or *9 of* adulthood. They were immobilised in 5 mM levamisole (Sigma-Aldrich) and mounted on 2% agarose pads. Neurons were visualized with a Zeiss Axio Imager M2 microscope. The software used was Zen Pro 2012. At least 30 worms were counted per condition.

#### Drug screening

##### Liquid culture

The animals were prepared for liquid culture as described above. Measurement was done using a Microtracker (Phylumtech) with standard parameters for *C. elegans*. All compounds were tested at 20 µM.

##### Solid Media

Worms were exposed from the L4 stage to compounds at 20 μM (with a 1% DMSO concentration) incorporated into NGM solid medium, or to NGM solid medium only as a control. All plates were streaked with OP50 E. coli. Briefly, 20 to 40 worms were picked and plated on the corresponding NGM medium (20 to 40 worms per plate for each condition, and each condition was done in triplicate) to complete observations of paralysis and neurodegeneration.

### Experimental design and statistical analysis

All experiments were conducted in triplicate, and the experimenters were blinded to the specimen genotype whenever possible (e.g., body size and motor neuron measurements). Quantitative data were presented as mean ± SEM, box plots, or Kaplan–Meier survival curves. GraphPad Prism v10.3.1 software was used for all statistical analyses. Statistical tests and sample sizes (*n*) are specified in the corresponding figure legends. For all datasets, outliers were eliminated using the ROUT method with a Q = 1%, and analyses were performed on the dataset without the outliers.

## Authorship contribution statement

A.E., P.D. and J.A.P. designed research; A.E., C.L., M.T. and C.M. performed experiments; A.E., C.L., M.T. and C.M. analyzed data; M.L. provided technical assistance in generating the KO line. A.E. and J.A.P. wrote the manuscript.

## Data availability

All data from this study are available from the corresponding author upon request

## Acknowledgments

This work was supported by the facilities of the Centre hospitalier de l’Université de Montréal (CHUM). We also thank the members of the Parker Lab for their support.

## Supporting information

### Supplementary Figures

**S1 Fig.**
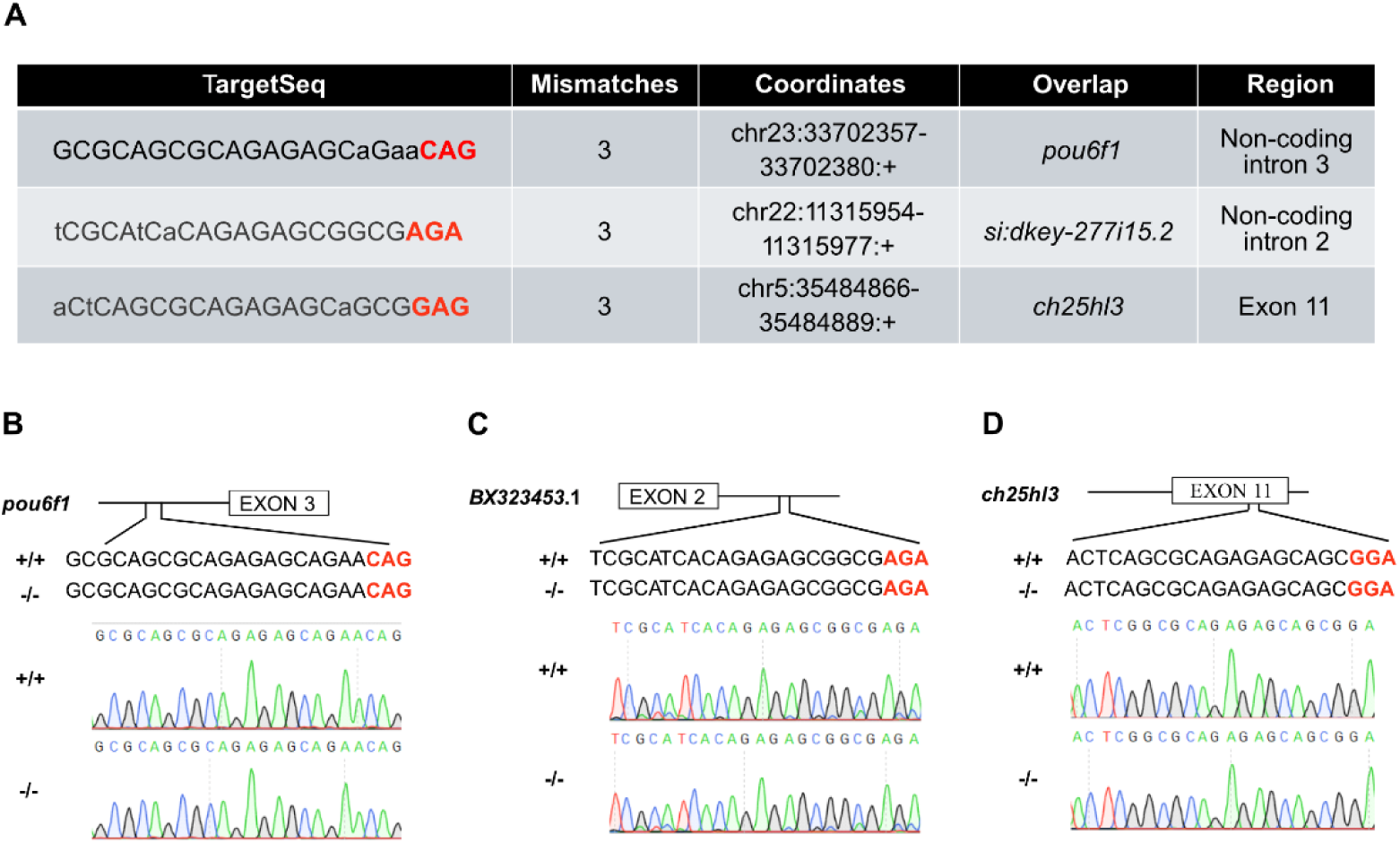
No evidence of off-target indels at the three most likely target sites for our *C9orf72* KO line. **A** Table listing the predicted off-target sequences, the number of mismatches with the original gRNA, chromosomal coordinates, the overlapping zebrafish gene, and the corresponding genomic region for the three most likely off-target sites, as predicted by CRISPRoff. The uppercase letters indicate matching base pairs and the lower cases differing pairs for the target sequences. **B-D** *C9orf72^-/-^* specimens are genetically identical to *C9orf72^+/+^* controls at the site of the possible off-target cutting predicted in the gene *pou6f1* (**B**), *si:dkey*-277i15.2 (**C**), and *ch25hl3* (**D**) for our gRNA.

**S2 Fig.**
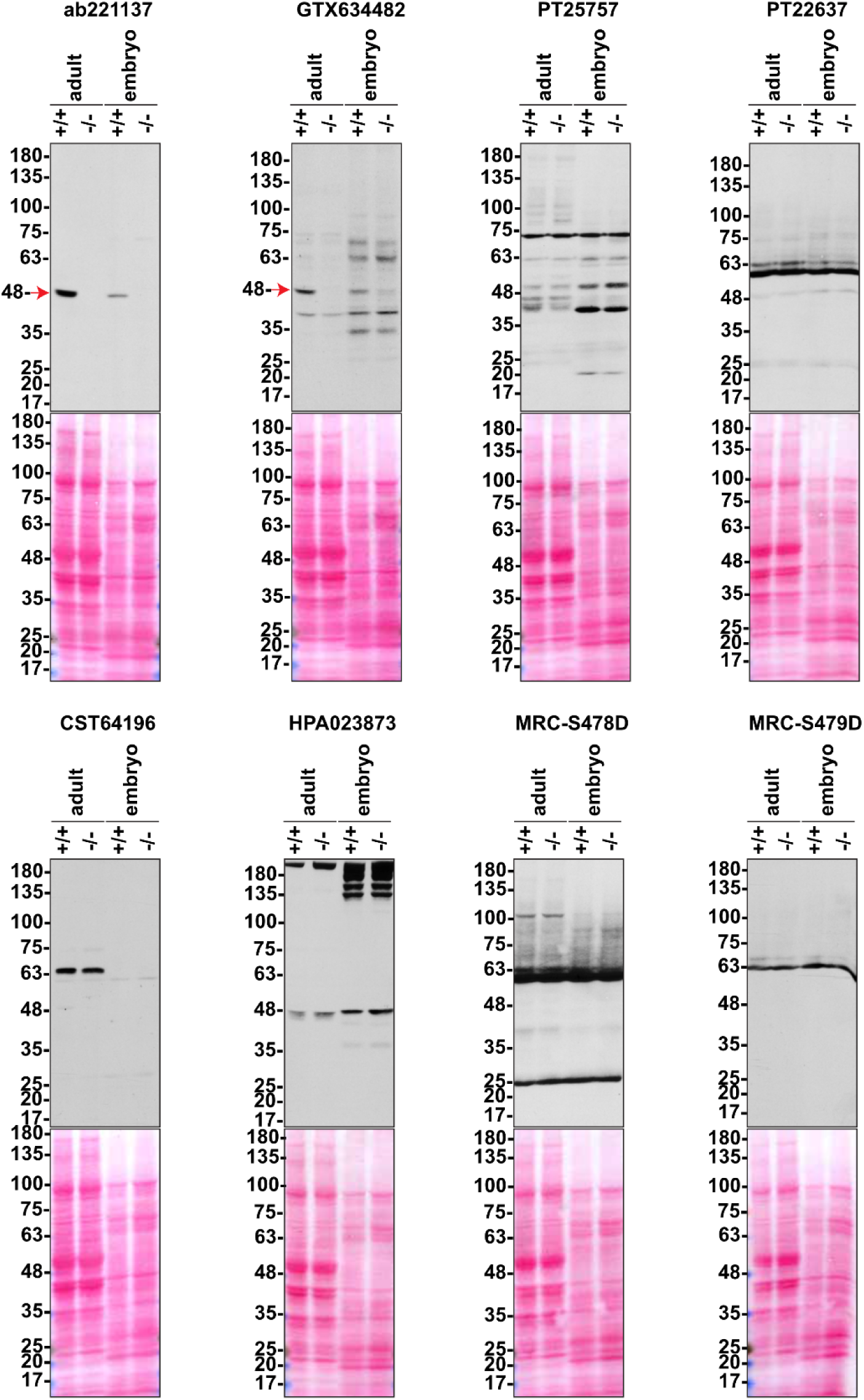
Analysis of eight antibodies known to recognize both human and murine C9orf72 for their ability to detect zebrafish C9orf72. Brain lysates from wild-type (+/+) and C9orf72 KO (-/-) adult zebrafish, as well as whole-larvae lysates at 2 dpf, were prepared and processed for immunoblotting with the indicated C9orf72 antibodies. The red arrows point to positive C9orf72 signals.

**S3 Fig.**
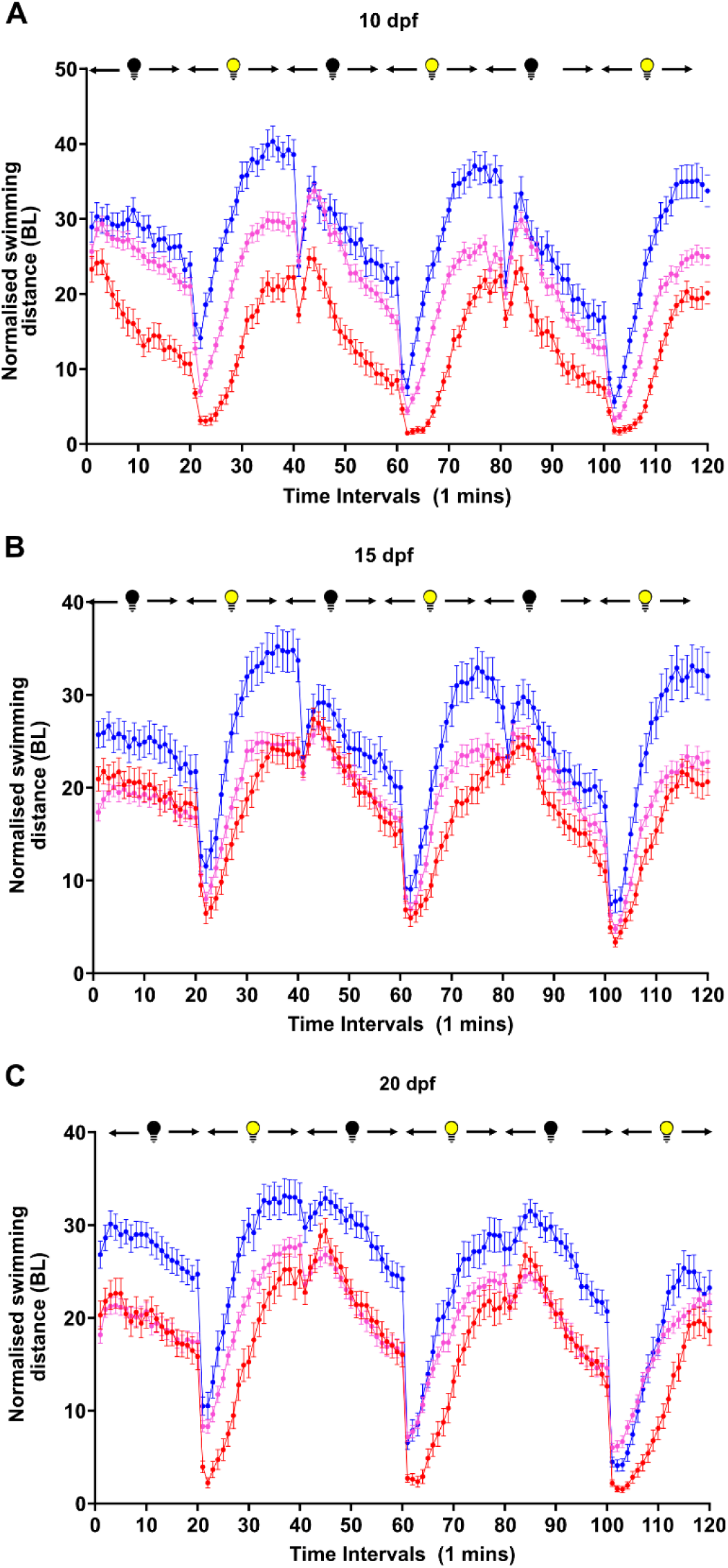
Highly stereotyped and conserved motor activity under alternating dark and light conditions across time and genotype in zebrafish larvae. **A-C** Point and connecting line with an error bar graph of the total swimming activity per minute normalized to body length observed with our phasic 120-minute dark-light program for *C9orf72^+/+^*, *C9orf72^-/+^* and *C9orf72^-/-^* 10 (**A**), 15 (**B**) and 20 (**C**) day post fertilization (dpf) larvae. N = 3, n = 72). Data are presented as mean ± SEM.

**S4 Fig.**
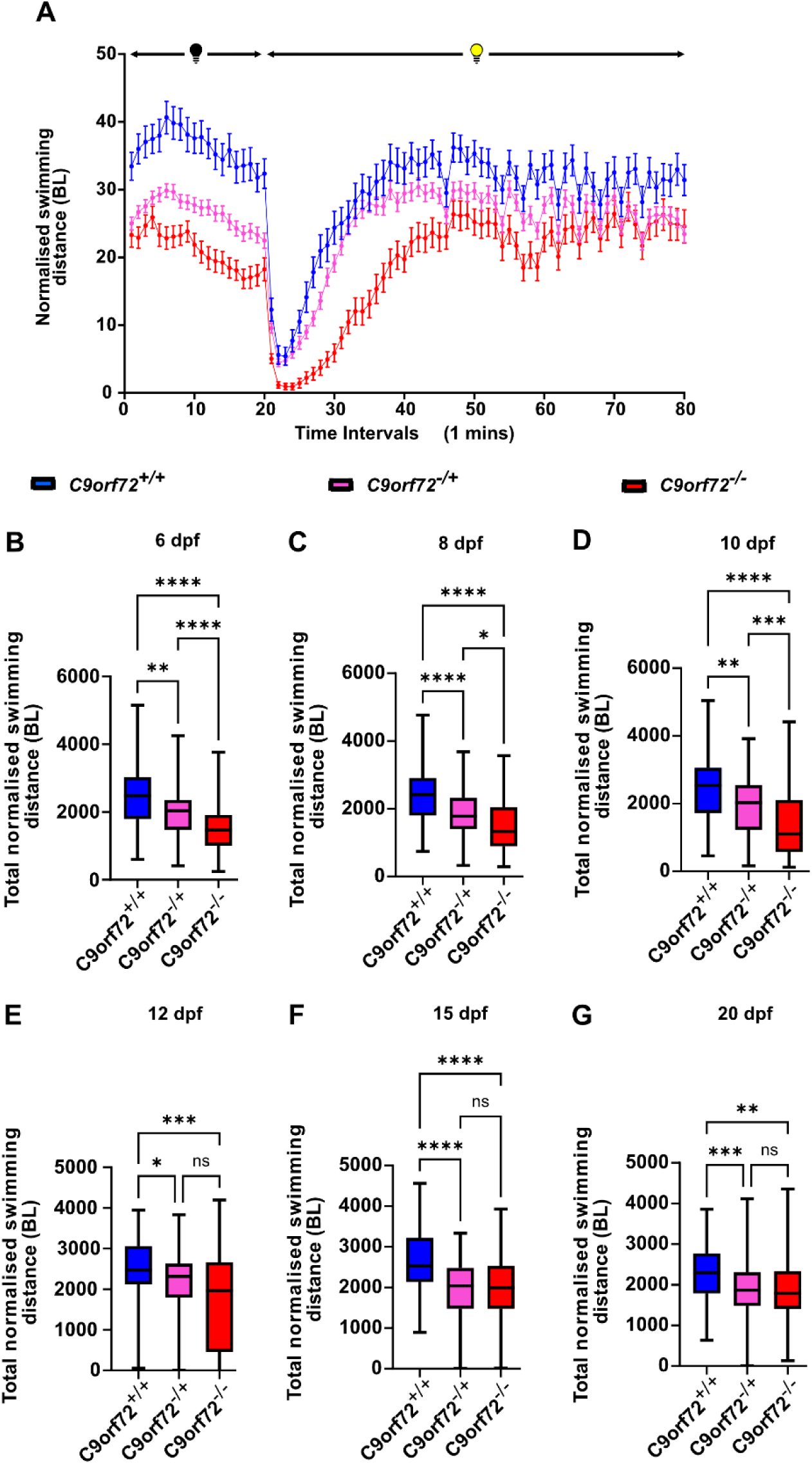
The knockout of *C9orf72* results in a persistent spontaneous swimming activity reduction in zebrafish larvae under a single Dark-light shift program. **(A)** Typical swimming distance normalized by average body length (BL) per minute pattern observed with our Dark (20 mins) / light (60 mins) program for *C9orf72^+/+^*, *C9orf72^-/+^* and *C9orf72^-/-^* 6 days post fertilization (dpf) larvae. Data are presented as mean ± SEM. (**B-G)** Quantitative analysis of total swimming distance normalized by average BL of larvae at 6, 8, 10, 12, 15 and 20 dpf during the SWAP period. (B-D) There is a significant deficit in normalized swimming activity for C9orf72^-/+^ and *C9orf72^-/-^* compared to *C9orf72^+/+^* controls and for *C9orf72^-/-^* compared to *C9orf72^+/-^* specimens at 6, 8 and 10 dpf. (**E**) *C9orf72^-/-^* and *C9orf72^-/+^* specimens show a significant reduction in normalized swimming compared to *C9orf72^+/+^* controls at 12, 15 and 20 dpf. Statistical tests: Kruskal-Walli test with Dunn’s multiple comparisons post-hoc test (**B-G**), ordinary one-way ANOVA with Tukey’s multiple comparisons post-hoc test (**H**). **** *p < 0.0001*, *** *p ≤ 0.001*, ** *p ≤ 0.01*, * *p ≤ 0.05* and NS *p > 0.05*. Boxplot extremities indicate maximum and minimum value, box limits indicate the range of the central 50% of the data, central line marks the median value. N = 3, n = 72 for each genotype except for *C9orf72^-/+,^* where n =144. N represents the number of experimental repeats from different clutches, and n represents the total number of larvae per genotype considered for the assay.

**S5 Fig.**
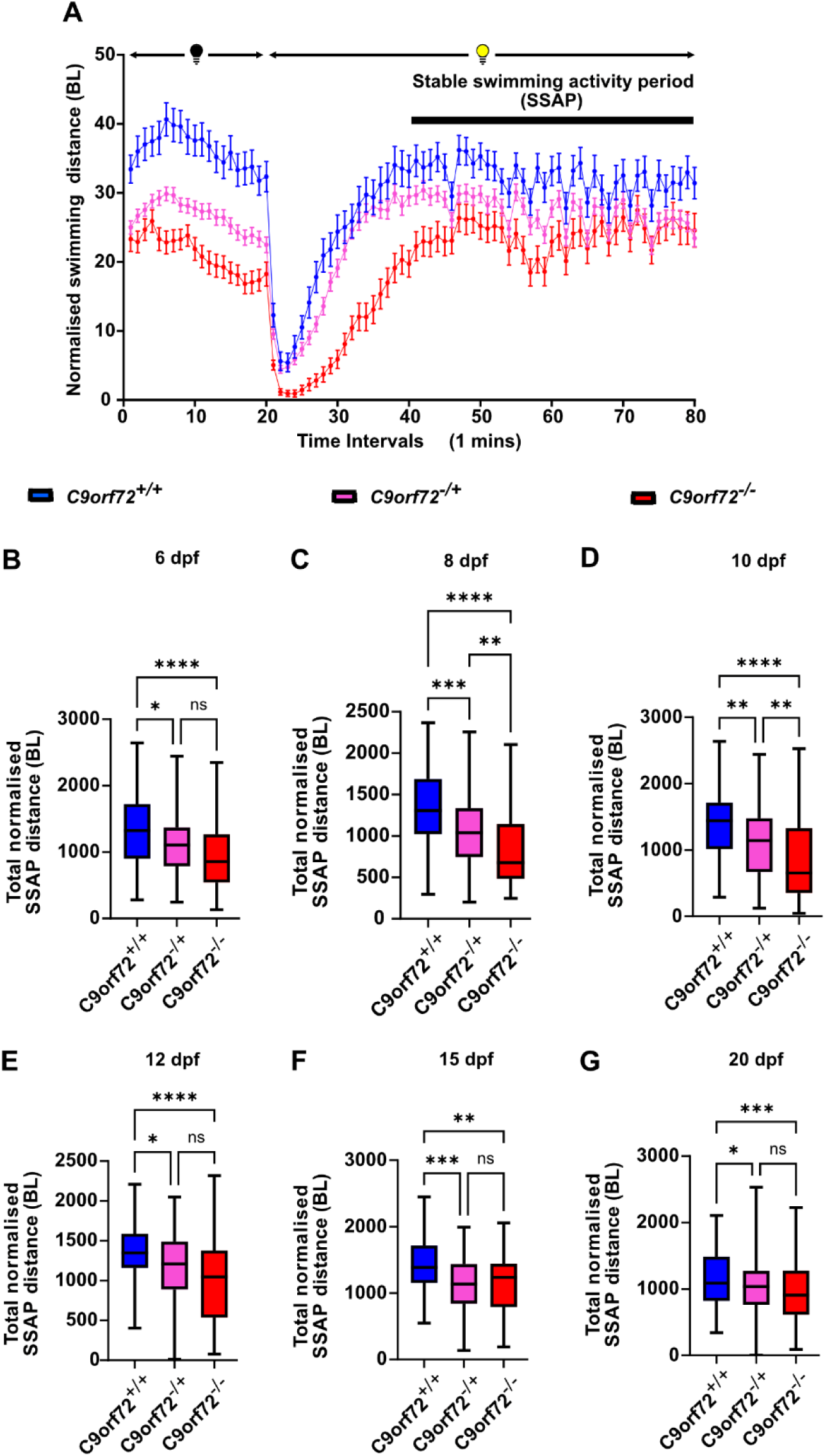
The knockout of *C9orf72* results in a persistent spontaneous swimming activity reduction in zebrafish larvae under stable light conditions. **(A)** Typical swimming distance normalized by average body length (BL) per minute pattern observed with our Dark (20 mins) / light (60 mins) program for *C9orf72^+/+^*, *C9orf72^-/+^* and *C9orf72^-/-^* 6 days post fertilization (dpf) larvae. The stable swimming activity period (SWAP) is defined as the period of relatively consistent swimming activity for each genotype, occurring approximately 20 minutes after the dark-to-light transition (minutes 40 to 80). Data are presented as mean ± SEM. (**B-G)** Quantitative analysis of total swimming distance normalized by average BL of larvae at 6, 8, 10, 12, 15 and 20 dpf during the SWAP period. (**B**) *C9orf72^-/-^ and C9orf72^-/+^* specimens show a significant reduction in normalized swimming compared to C9orf72^+/+^ controls at 6 dpf. (**C-D**) There is a significant deficit in normalized swimming activity for C9orf72^-/+^ and *C9orf72^-/-^* compared to *C9orf72^+/+^* controls and for *C9orf72^-/-^* compared to *C9orf72^+/-^* specimens at 8 and 10 dpf. (**E, F, G**) *C9orf72^-/-^* and *C9orf72^-/+^* specimens show a significant reduction in normalized swimming compared to *C9orf72^+/+^* controls at 12, 15 and 20 dpf. Statistical tests: ordinary one-way ANOVA with Tukey’s multiple comparisons post-hoc test (**A**, **H**) Kruskal-Walli test with Dunn’s multiple comparisons post-hoc test (**C-G**), Welch and Brown-Forsythe ANOVA test with Dunnett’s T3 multiple comparisons post-hoc test **** *p < 0.0001*, *** *p ≤ 0.001*, ** *p ≤ 0.01*, * *p ≤ 0.05* and NS *p > 0.05*. Boxplot extremities indicate maximum and minimum value, box limits indicate the range of the central 50% of the data, central line marks the median value. N = 3, n = 72 for each genotype except for *C9orf72^-/+,^* where n =144. N represents the number of experimental repeats from different clutches, and n represents the total number of larvae per genotype considered for the assay.

**S6 Fig.**
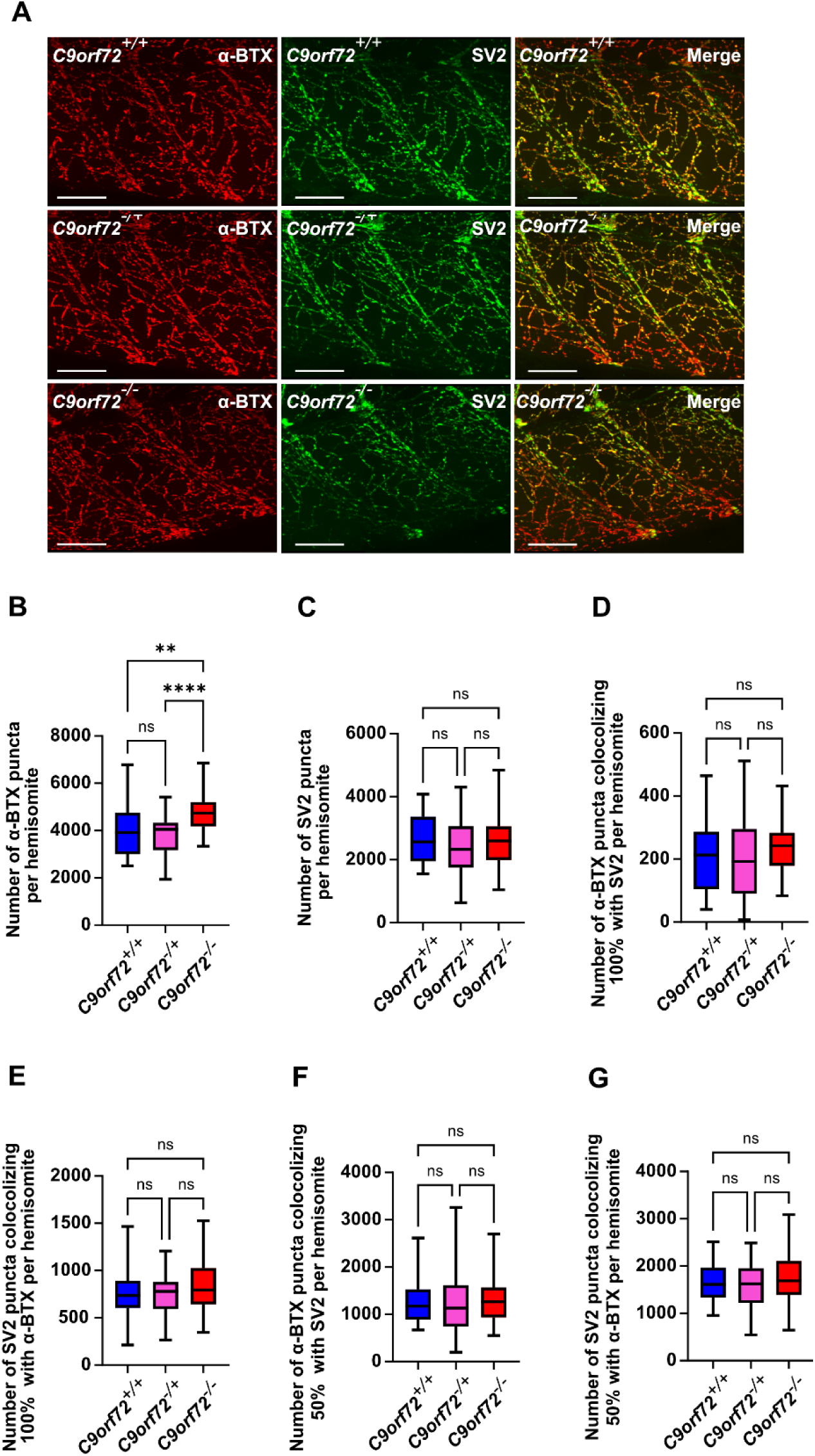
Complete loss of C9orf72 induces a mild postsynaptic alteration at the neuromuscular junction in zebrafish larvae. **(A)** Representative confocal images of co-immunostaining at 6 days post-fertilization (dpf) for presynaptic (SV2, green) and postsynaptic (α-bungarotoxin, red) markers in hemisegment NMJs across all three C9orf72 related genotypes. **(B)** Quantitative analysis reveals a significant increase in the total number of α-bungarotoxin (α-BTX) labeled postsynaptic puncta in *C9orf72^-/-^* larvae compared to *C9orf72^-/+^* and *C9orf72^+/+^* controls. **(C)** No significant differences were observed in the total number of presynaptic puncta (SV2) across genotypes. **(D-E)** No significant differences were detected in the number of postsynaptic puncta (α-BTX) that fully colocalize (100%) with presynaptic SV2 or in the number of SV2 puncta fully colocalizing (100%) with α-BTX. **(F-G)** Similarly, no significant differences were observed in partial colocalization (50%) between α-BTX and SV2a puncta. Statistical tests : Ordinary one-way ANOVA with Tukey’s multiple comparisons post-hoc test **(B,E,G)**, Kruskal-Wallis test with test Dunn’s multiple comparisons post-hoc test **(C,F)**, Welch and Brown-Forsythe ANOVA with Dunnett’s T3 multiple comparisons test **(D).** *** *p ≤ 0.001* and NS *p > 0.05*. Boxplot extremities indicate maximum and minimum values; box limits represent the interquartile range (central 50%), and the central line marks the median value. N = 12 (total number of distinct specimens); n = 34–36 (total number of hemisegments analyzed per genotype). Scale bars = 50 µm.

**S7 Fig.**
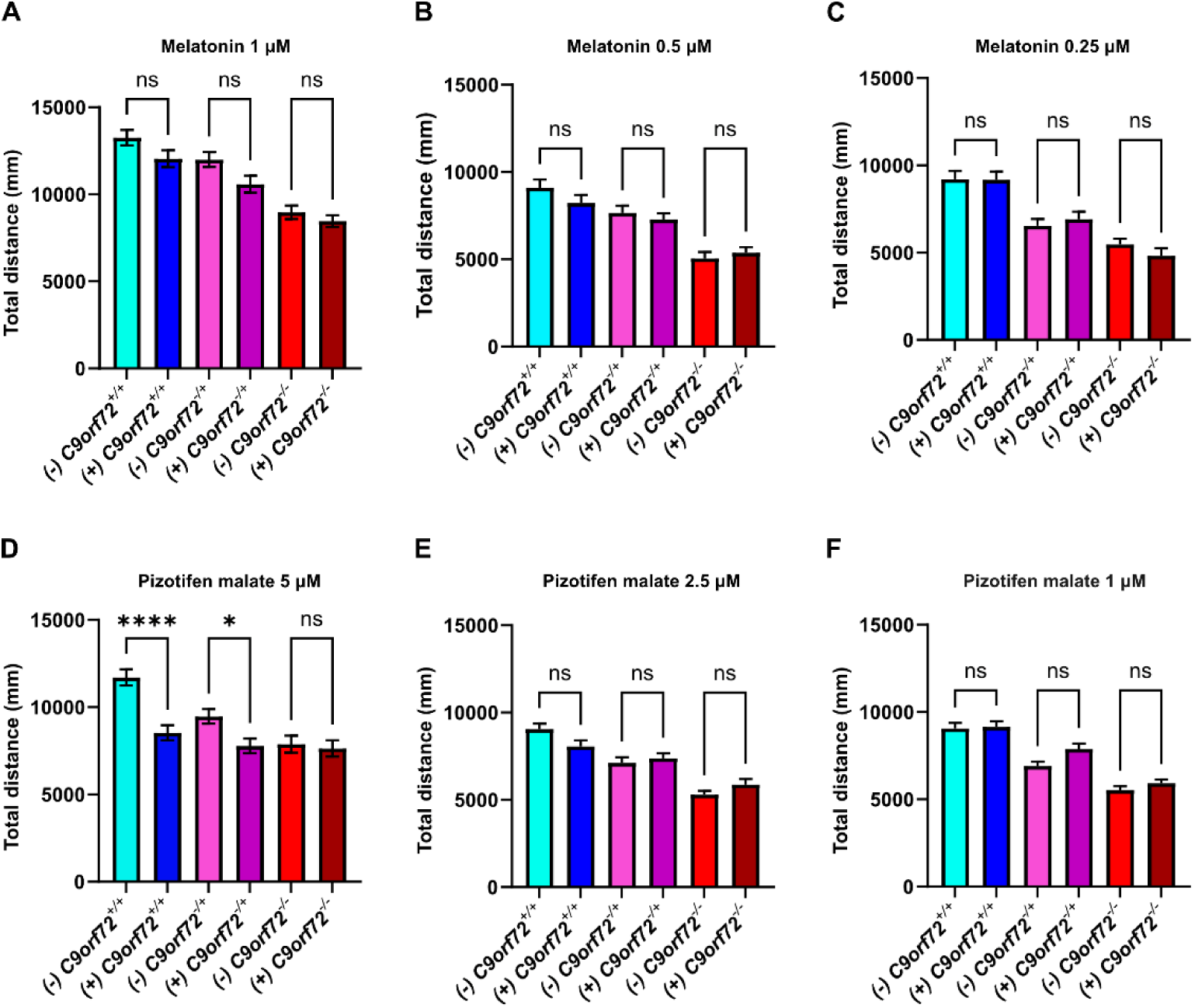
Effect of melatonin and pizotifen malate on zebrafish larvae swimming activity of all three genotypes at different concentrations. Melatonin treatment did not improve the swimming deficit in *C9orf72* KO larvae at any tested concentration, nor did higher concentrations of PM. Bar graphs represent the total swimming distance of 6 dpf zebrafish subjected to the phasic light-dark paradigm. **(A-C)** Melatonin treatment had no significant effect on total swimming activity across all genotypes and concentrations tested. **(D-F)** Higher doses of PM significantly reduced swimming activity in *C9orf72^+/+^* and *C9orf72^+/-^* larvae, suggesting potential toxicity **(D)**, whereas no significant effect was observed for any genotype at other tested doses **(E-F).** Statistical tests: Kruskal-Wallis test with test Dunn’s multiple comparisons post-hoc test **(B-F),** Welch and Brown-Forsythe ANOVA with Dunnett’s T3 multiple comparisons test **(A**). **** *p < 0.0001*, ** *p ≤ 0.01*, NS *p > 0.05*. (-) indicates untreated specimens, (+) indicates treated specimens. Sample size: N = 2-4 (number of independent swimming assays from different clutches); n = 32-80 (total number of individual specimens). Data are presented as mean ± SEM.

### Supplementary Table

**S1 Table.**
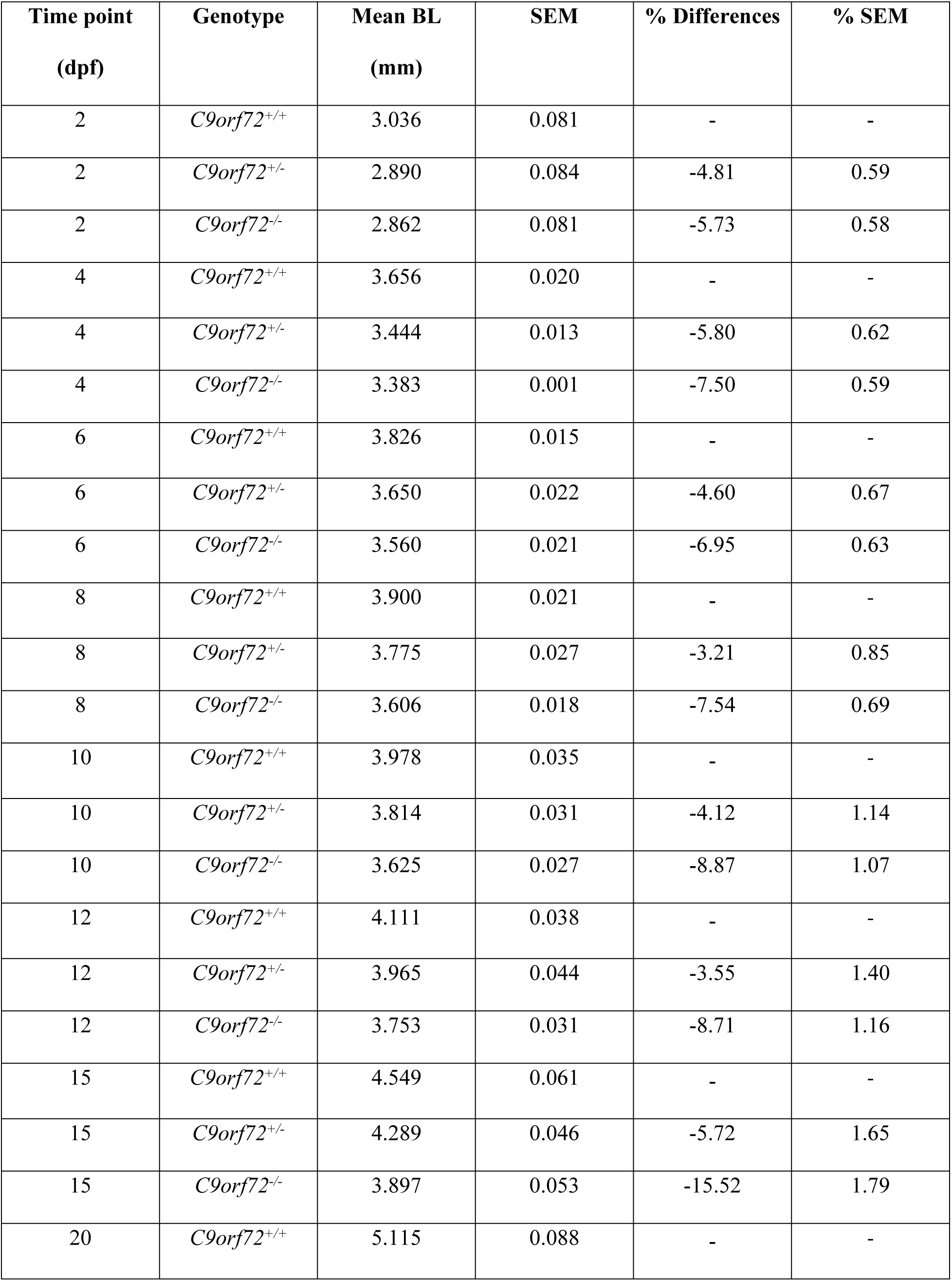
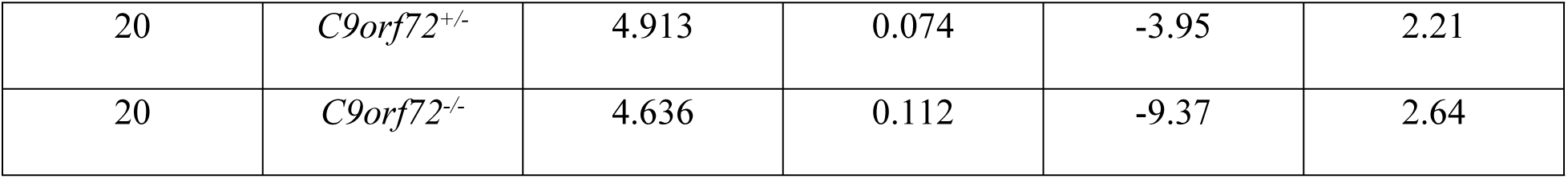
Body length averages and relevant statistics for zebrafish larvae of all three genotypes of the C9orf72 knockout. Table of all the results and associated statistics concerning the measurement of body length (BL) in millimeters (mm) of *C9orf72^+/+^*, *C9orf72^-/^*^+^, and *C9orf72^-/-^* specimens for all the time points considered. Days post-fertilisation (dpf); standard error mean (SEM).

| Time point<br>(dpf) | Genotype | Mean BL<br>(mm) | SEM | % Differences | % SEM |
| --- | --- | --- | --- | --- | --- |
| 2 | *C9orf72*<sup>+/+</sup> | 3.036 | 0.081 | - | - |
| 2 | *C9orf72*<sup>+/-</sup> | 2.890 | 0.084 | -4.81 | 0.59 |
| 2 | *C9orf72*<sup>-/-</sup> | 2.862 | 0.081 | -5.73 | 0.58 |
| 4 | *C9orf72*<sup>+/+</sup> | 3.656 | 0.020 | - | - |
| 4 | *C9orf72*<sup>+/-</sup> | 3.444 | 0.013 | -5.80 | 0.62 |
| 4 | *C9orf72*<sup>-/-</sup> | 3.383 | 0.001 | -7.50 | 0.59 |
| 6 | *C9orf72*<sup>+/+</sup> | 3.826 | 0.015 | - | - |
| 6 | *C9orf72*<sup>+/-</sup> | 3.650 | 0.022 | -4.60 | 0.67 |
| 6 | *C9orf72*<sup>-/-</sup> | 3.560 | 0.021 | -6.95 | 0.63 |
| 8 | *C9orf72*<sup>+/+</sup> | 3.900 | 0.021 | - | - |
| 8 | *C9orf72*<sup>+/-</sup> | 3.775 | 0.027 | -3.21 | 0.85 |
| 8 | *C9orf72*<sup>-/-</sup> | 3.606 | 0.018 | -7.54 | 0.69 |
| 10 | *C9orf72*<sup>+/+</sup> | 3.978 | 0.035 | - | - |
| 10 | *C9orf72*<sup>+/-</sup> | 3.814 | 0.031 | -4.12 | 1.14 |
| 10 | *C9orf72*<sup>-/-</sup> | 3.625 | 0.027 | -8.87 | 1.07 |
| 12 | *C9orf72*<sup>+/+</sup> | 4.111 | 0.038 | - | - |
| 12 | *C9orf72*<sup>+/-</sup> | 3.965 | 0.044 | -3.55 | 1.40 |
| 12 | *C9orf72*<sup>-/-</sup> | 3.753 | 0.031 | -8.71 | 1.16 |
| 15 | *C9orf72*<sup>+/+</sup> | 4.549 | 0.061 | - | - |
| 15 | *C9orf72*<sup>+/-</sup> | 4.289 | 0.046 | -5.72 | 1.65 |
| 15 | *C9orf72*<sup>-/-</sup> | 3.897 | 0.053 | -15.52 | 1.79 |
| 20 | *C9orf72*<sup>+/+</sup> | 5.115 | 0.088 | - | - |
| 20 | *C9orf72*<sup>+/-</sup> | 4.913 | 0.074 | -3.95 | 2.21 |
| 20 | *C9orf72*<sup>-/-</sup> | 4.636 | 0.112 | -9.37 | 2.64 |

**S2 Table.**
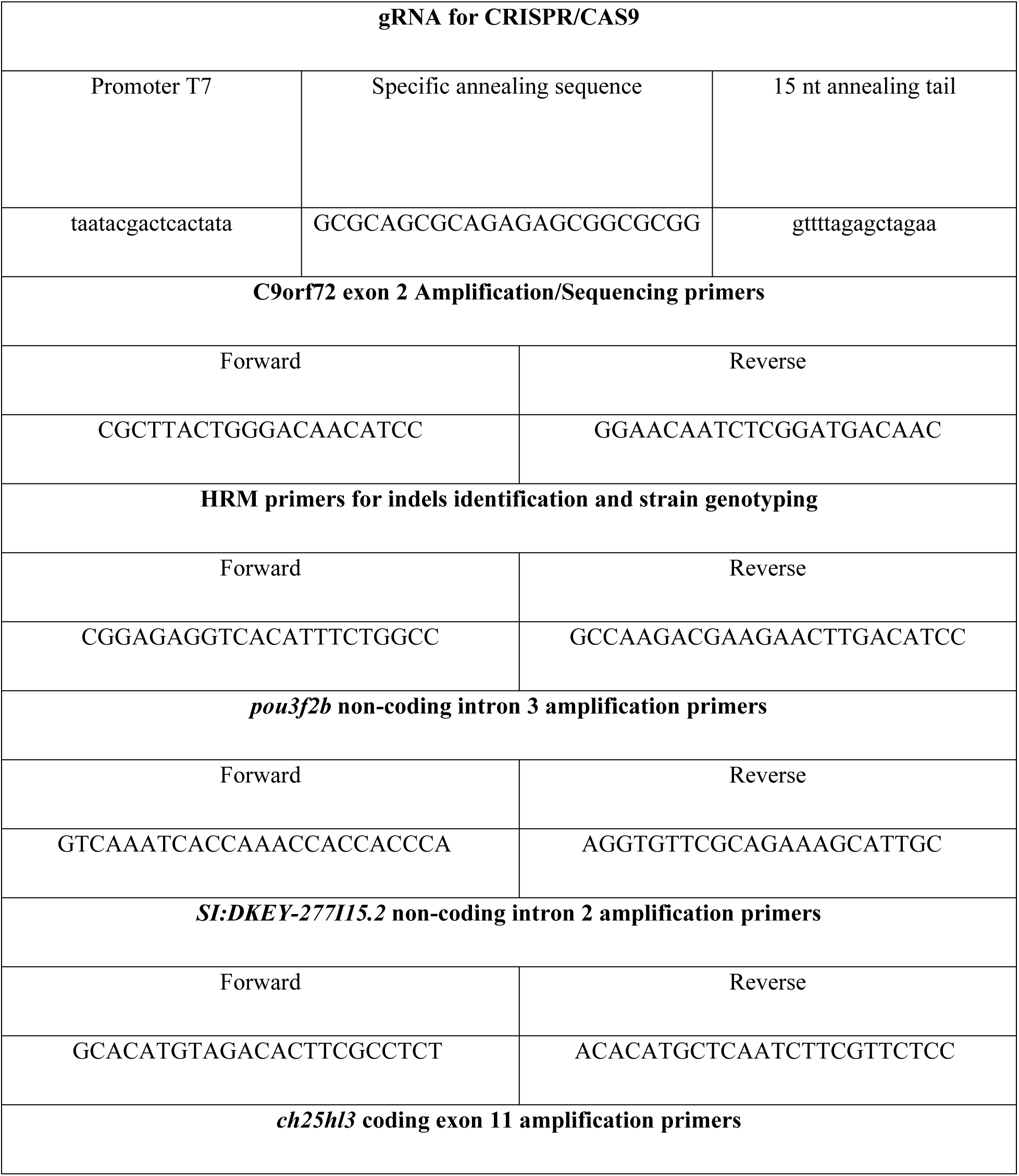
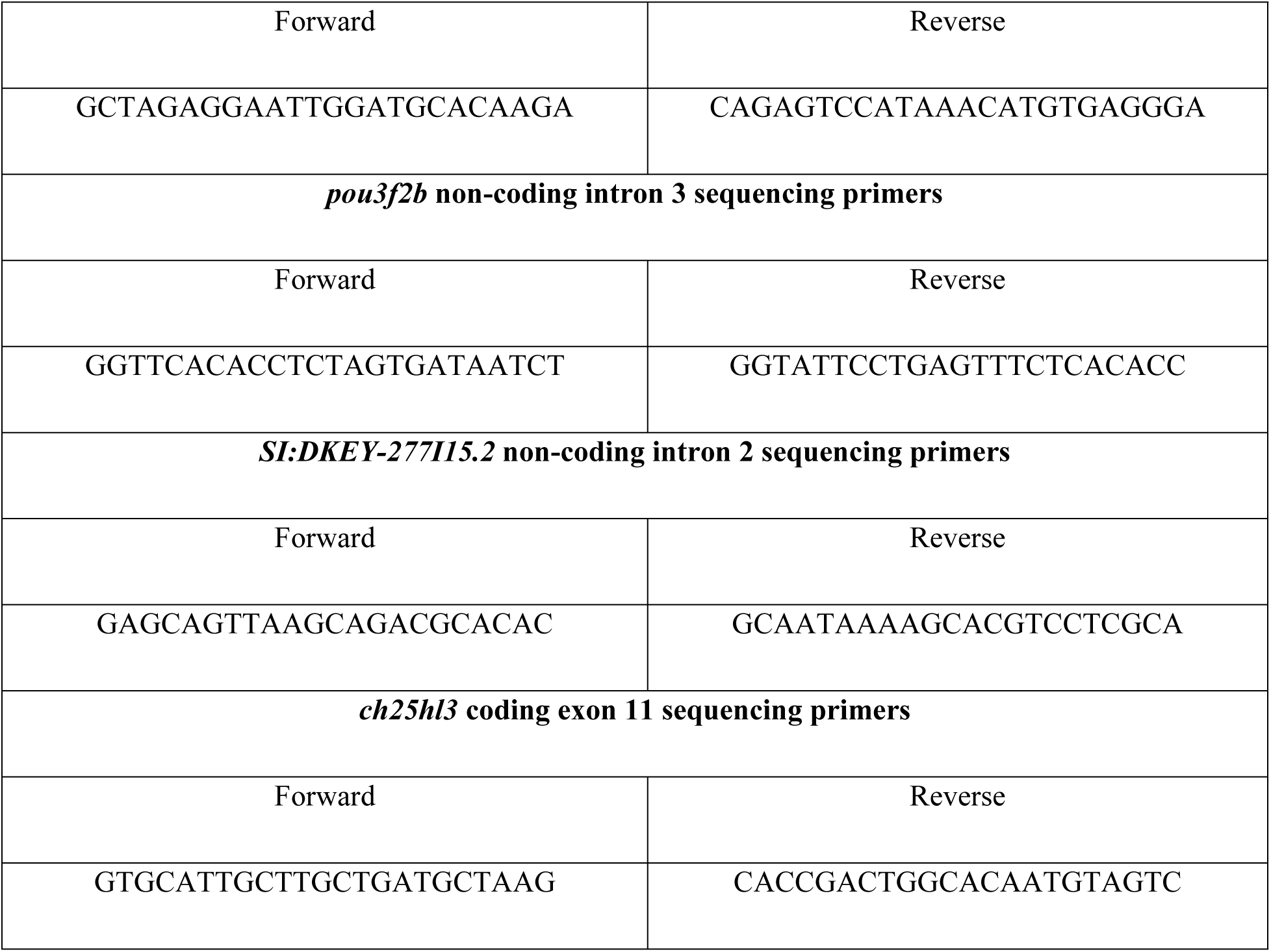
Oligonucleotide sequences. Table of all the oligonucleotide sequences that were used for the generation of the C9orf72 LOF line by CRISPR/Cas9, PCR amplification, HRM, and sequencing.

**S3 Table.**
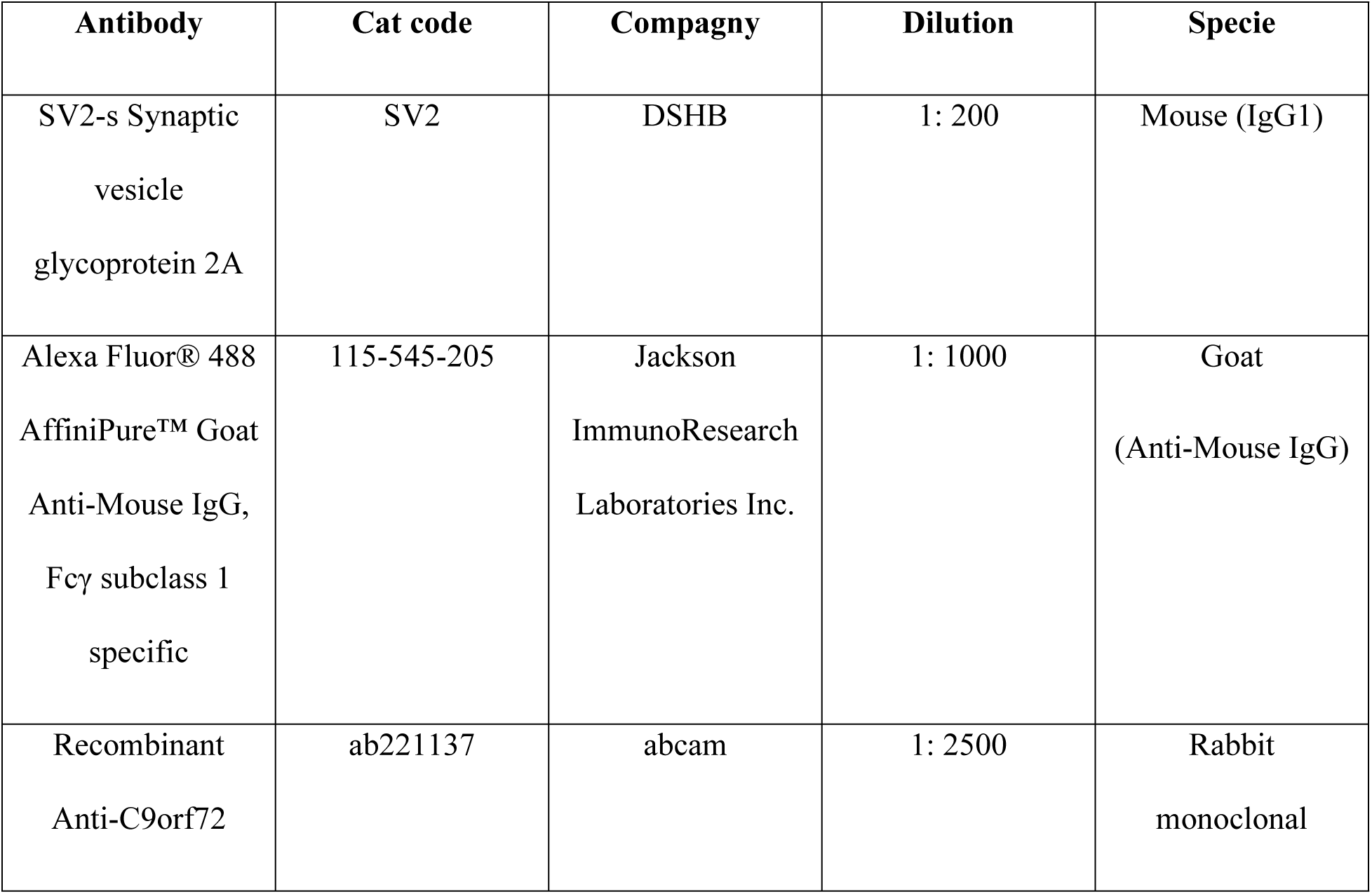
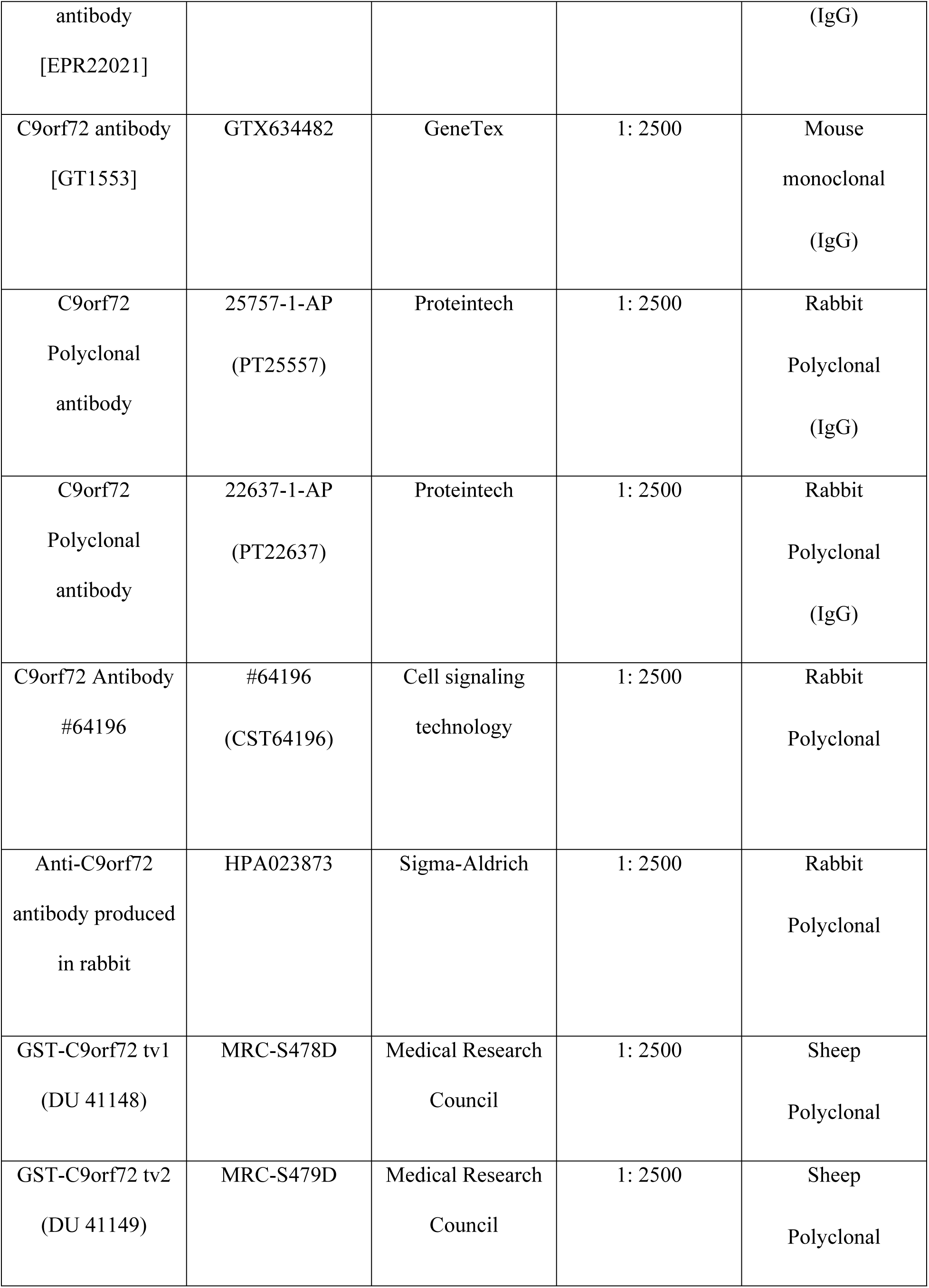
Antibodies. . Table of all the antibodies used for the immunofluorescence assay (neuromuscular junction imaging) and the Western blots (C9orf72 detection).

**S4 Table.**
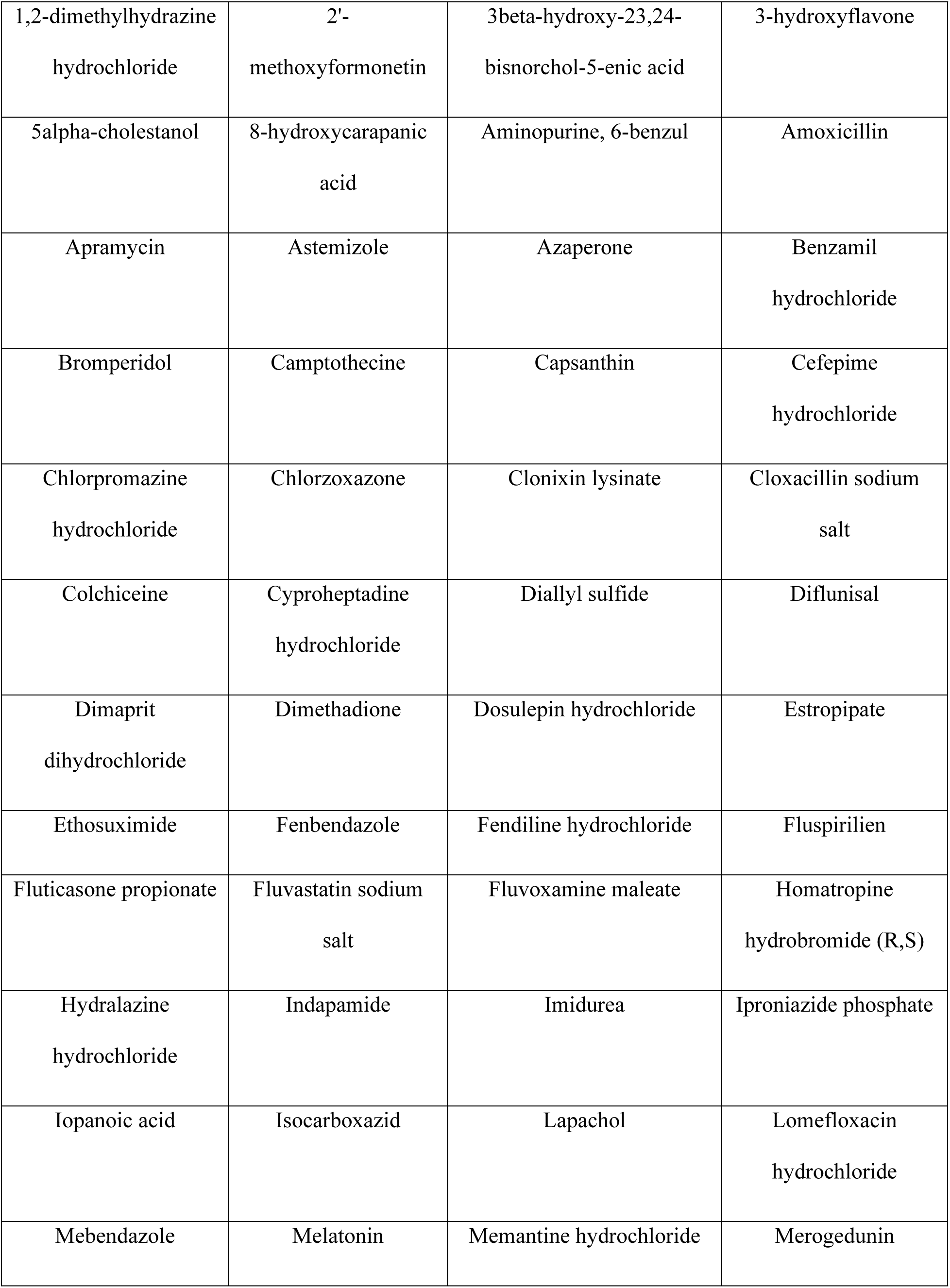
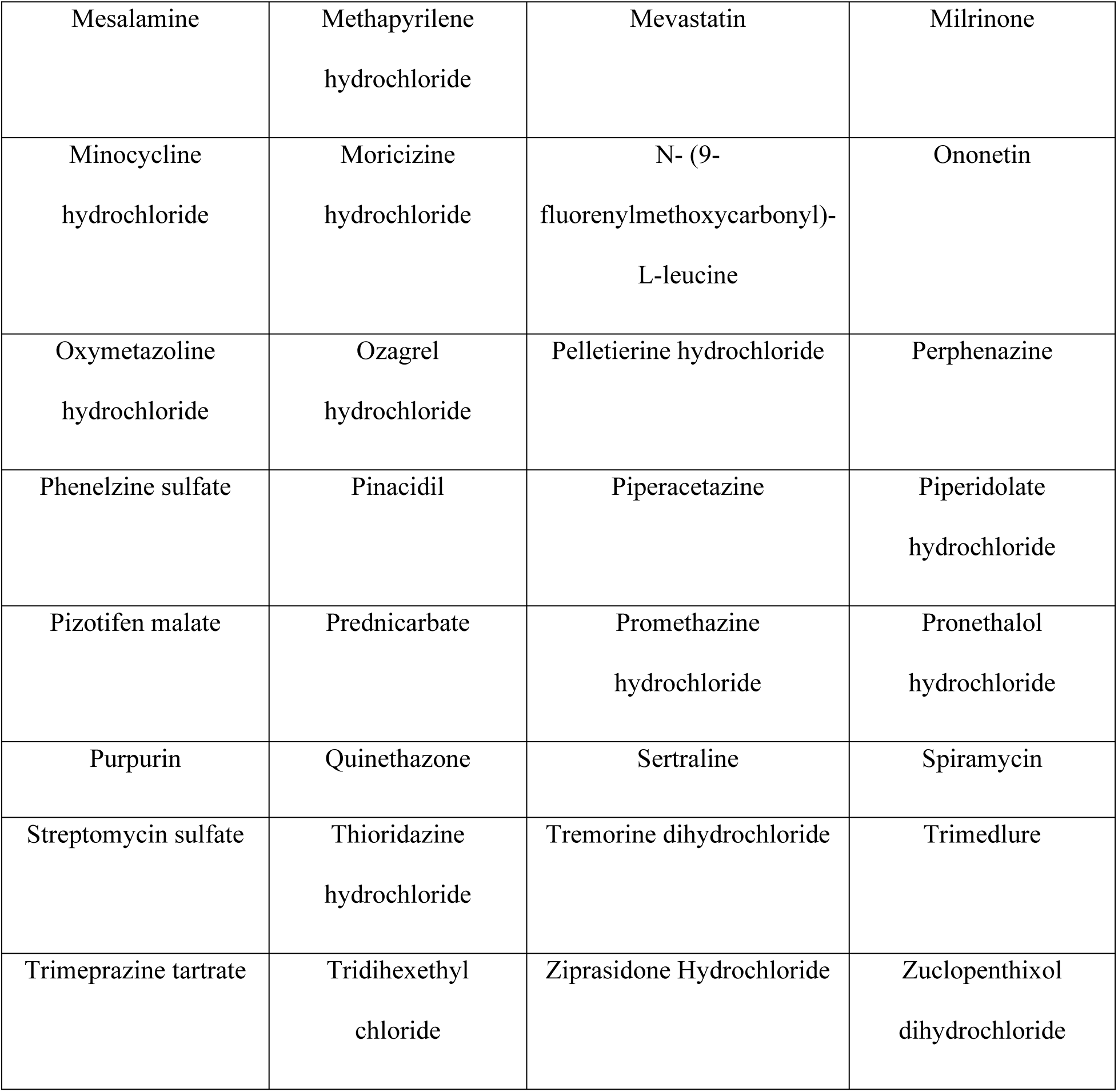
Molecules identified that ameliorate *alfa-1(ok3063)* motility defect in liquid culture. Table of the 80 compounds identified that were able to ameliorate the motility defect of *alfa-1(ok3063) C*. elegans animals in liquid culture.

**S5 table.** Molecules identified that improve *alfa-1 (ok3062)* age-dependent paralysis and neurodegeneration phenotypes. Table of the molecules identified that improve the age-dependent paralysis and neurodegeneration phenotypes of *alfa-1 (ok3062) C. elegans* animals on solid media. Statistical tests and parameters are the same as described for melatonin and PM in supplementary data.

| **Compounds** | **Paralysis curve comparison**<br>**with untreated, p value** | **Neurodegeneration %**<br>**comparison with untreated,**<br>**p value** |
| --- | --- | --- |
| Dimethatione | 0.0002 | 0.0028 |
| Fluvoxamine maleate | 0.0026 | 0.0045 |
| Melatonin | $\leq 0.0001$ | 0.0067 |
| Mesalamine | 0.0101 | 0.0028 |
| Milirinone | 0.0146 | 0.002 |
| Perphenazine | 0.1141 | 0.0045 |
| Pizotifen malate | $\leq 0.0001$ | 0.0281 |
| Prednicarbate | 0.0249 | 0.001 |
| Thioridazine hydrochloride | $\leq 0.0001$ | 0.0045 |
| Tremorine dihydrochloride | 0.0219 | 0.0045 |
| Tridihexethyl chloride | 0.0472 | 0.0045 |
| Zuclopenthixol dihydrochloride | 0.0159 | 0.0045 |

## Notes

### Competing Interest Statement

The authors have declared no competing interest.

## References

1. Mead RJ, Shan N, Reiser HJ, Marshall F, Shaw PJ. Amyotrophic lateral sclerosis: a neurodegenerative disorder poised for successful therapeutic translation. Nat Rev Drug Discov. 2023;22(3):185–212. doi: 10.1038/s41573-022-00612-2.

2. DeJesus-Hernandez M, Mackenzie IR, Boeve BF, Boxer AL, Baker M, Rutherford NJ, et al. Expanded GGGGCC hexanucleotide repeat in noncoding region of C9ORF72 causes chromosome 9p-linked FTD and ALS. Neuron. 2011;72(2):245–56. doi: 10.1016/j.neuron.2011.09.011.

3. Renton AE, Majounie E, Waite A, Simón-Sánchez J, Rollinson S, Gibbs JR, et al. A hexanucleotide repeat expansion in C9ORF72 is the cause of chromosome 9p21-linked ALS-FTD. Neuron. 2011 Oct 20;72(2):257–68. doi: 10.1016/j.neuron.2011.09.010.

4. Majounie E, Renton AE, Mok K, Dopper EGP, Waite A, Rollinson S, et al. Frequency of the C9orf72 hexanucleotide repeat expansion in patients with amyotrophic lateral sclerosis and frontotemporal dementia: a cross-sectional study. Lancet Neurol. 2012 Apr;11(4):323–30. doi: 10.1016/S1474-4422(12)70043-1.

5. Smeyers J, Banchi EG, Latouche M. C9ORF72: what it is, what it does, and why it matters. Front Cell Neurosci. 2021 May 5;15:661447. doi: 10.3389/fncel.2021.661447.

6. Waite AJ, Bäumer D, East S, Neal J, Morris HR, Ansorge O, et al. Reduced C9orf72 protein levels in frontal cortex of amyotrophic lateral sclerosis and frontotemporal degeneration brain with the C9orf72 hexanucleotide repeat expansion. Neurobiol Aging. 2014;35(8):1779.e5–13. doi: 10.1016/j.neurobiolaging.2014.01.016.

7. Balendra R, Isaacs AM. C9orf72-mediated ALS and FTD: multiple pathways to disease. Nat Rev Neurol. 2018;14(9):544–58. doi: 10.1038/s41582-018-0047-2.

8. Burberry A, Wells MF, Lim RG, Coughlin M, Bu J, Marchetto MC, et al. Loss-of-function mutations in the C9ORF72 mouse ortholog cause fatal autoimmune disease. Sci Transl Med. 2016;8(347):347ra93. doi: 10.1126/scitranslmed.aaf6038.

9. O’Rourke JG, Bogdanik L, Yáñez A, Lall D, Wolf AJ, Muhammad AKMG, et al. C9orf72 is required for proper macrophage and microglial function in mice. Science. 2016;351(6279):1324–9. doi: 10.1126/science.aaf1064.

10. Atanasio A, Decman V, White D, Ramos M, Ikiz B, Lee HC, et al. C9orf72 ablation causes immune dysregulation characterized by leukocyte expansion, autoantibody production, and glomerulonephropathy in mice. Sci Rep. 2016;6:23204. doi: 10.1038/srep23204.

11. Lopez-Herdoiza MB, Bauché S, Wilmet B, Le Duigou C, Roussel D, Frah M, et al. C9ORF72 knockdown triggers FTD-like symptoms and cell pathology in mice. Front Cell Neurosci. 2023 Apr 17;17:1155929. doi: 10.3389/fncel.2023.1155929.

12. Chia K, Klingseisen A, Sieger D, Priller J. Zebrafish as a model organism for neurodegenerative disease. Front Mol Neurosci. 2022;15:940484. doi: 10.3389/fphar.2021.713963.

13. Paquet D, Bhat R, Sydow A, Mandelkow EM, Berg S, Hellberg S, et al. A zebrafish model of tauopathy allows in vivo imaging of neuronal cell death and drug evaluation. J Clin Invest. 2009;119(5):1382–95. doi: 10.1172/JCI37537DS1.

14. Flinn LJ, Keatinge M, Bretaud S, Mortiboys H, Matsui H, De Felice E, et al. TigarB causes mitochondrial dysfunction and neuronal loss in PINK1 deficiency. Ann Neurol. 2013;74(6):837–47. doi: 10.1136/jnnp-2013-306573.16.

15. Keatinge M, Bui H, Menke A, Chen YC, Sokol AM, Bai Q, et al. Glucocerebrosidase 1-deficient Danio rerio mirror key pathological aspects of human Gaucher disease and provide evidence of early microglial activation preceding alpha-synuclein-independent neuronal cell death. Hum Mol Genet. 2015 Sep 16;24(23):6640–52. doi: 10.1093/hmg/ddv369.

16. Larbalestier H, Keatinge M, Watson L, White E, Gowda S, Wei W, et al. GCH1 deficiency activates brain innate immune response and impairs tyrosine hydroxylase homeostasis. J Neurosci. 2022;42(4):702–16. doi: 10.1523/jneurosci.0653-21.2021.

17. Ciura S, Lattante S, Le Ber I, Latouche M, Tostivint H, Brice A, et al. Loss of function of C9orf72 causes motor deficits in a zebrafish model of amyotrophic lateral sclerosis. Ann Neurol. 2013;74(2):180–90. doi: 10.1002/ana.23946.

18. Butti Z, Pan YE, Giacomotto J, Patten SA. Reduced C9orf72 function leads to defective synaptic vesicle release and neuromuscular dysfunction in zebrafish. Commun Biol. 2021;4:792. doi: 10.1038/s42003-021-02302-y.

19. Jaroszynska N, Salzinger A, Tsarouchas TM, Becker CG, Becker T, Lyons DA, et al. C9ORF72 deficiency results in degeneration of the zebrafish retina in vivo. J Neurosci. 2024;44(25):e2128232024. doi: 10.1523/JNEUROSCI.2128-23.2024.

20. Giacomotto J, Ségalat L. High-throughput screening and small animal models, where are we? Br J Pharmacol. 2010 May;160(2):204–16. doi: 10.1111/j.1476-5381.2010.00725.x.

21. Caldwell KA, Willicott CW, Caldwell GA. Modeling neurodegeneration in Caenorhabditis elegans. Dis Model Mech. 2020 Oct 26;13(10):dmm046110. doi: 10.1242/dmm.046110.

22. Roussos A, Kitopoulou K, Borbolis F, Palikaras K. Caenorhabditis elegans as a model system to study human neurodegenerative disorders. Biomolecules. 2023;13:478. doi: 10.3390/biom13030478.

23. Therrien M, Rouleau GA, Dion PA, Parker JA. Deletion of C9ORF72 results in motor neuron degeneration and stress sensitivity in C. elegans. PLoS One. 2013;8(12):e83450. doi: 10.1371/journal.pone.0083450.

24. Schmeisser K, Fardghassemi Y, Parker JA. A rapid chemical-genetic screen utilizing impaired movement phenotypes in C. elegans: input into genetics of neurodevelopmental disorders. Exp Neurol. 2017 Jul;293:101–14. doi: 10.1016/j.expneurol.2017.03.022.

25. Patten SA, Aggad D, Martinez J, Tremblay E, Petrillo J, Armstrong GA, et al. Neuroleptics as therapeutic compounds stabilizing neuromuscular transmission in amyotrophic lateral sclerosis. JCI Insight. 2017 Nov 16;2(22):e97152. doi: 10.1172/jci.insight.97152.

26. Doyle JJ, Maios C, Vrancx C, Duhaime S, Chitramuthu B, Bennett HPJ, et al. Chemical and genetic rescue of in vivo progranulin-deficient lysosomal and autophagic defects. Proc Natl Acad Sci U S A. 2021 Jun 22;118(25):e2022115118. doi: 10.1073/pnas.2022115118.

27. Tossing G, Livernoche R, Maios C, Bretonneau C, Labarre A, Parker JA. Genetic and pharmacological PARP inhibition reduces axonal degeneration in C. elegans models of ALS. Hum Mol Genet. 2022 Sep 29;31(19):3313–24. doi: 10.1093/hmg/ddac116.

28. Anthon C, Corsi GI, Gorodkin J. CRISPRon/off: CRISPR/Cas9 on- and off-target gRNA design. Bioinformatics. 2022;38(20):5437–9. doi: 10.1093/bioinformatics/btac697.

29. Lejeune F. Nonsense-mediated mRNA decay, a finely regulated mechanism. Biomedicines. 2022 Jan 10;10(1):141. doi: 10.3390/biomedicines10010141.

30. Laflamme C, McKeever PM, Kumar R, Schwartz J, Kolahdouzan M, Chen CX, et al. Implementation of an antibody characterization procedure and application to the major ALS/FTD disease gene C9ORF72. Elife. 2019;8:e48363. doi: 10.7554/eLife.48363.

31. Basnet RM, Zizioli D, Taweedet S, Finazzi D, Memo M. Zebrafish larvae as a behavioral model in neuropharmacology. Biomedicines. 2019;7(1):23. doi: 10.3390/biomedicines7010023.

32. Babin PJ, Goizet C, Raldúa D. Zebrafish models of human motor neuron diseases: advantages and limitations. Prog Neurobiol. 2014;118:36–58. doi: 10.1016/j.pneurobio.2014.03.001.

33. Gjorgjieva J, Biron D, Haspel G. Neurobiology of Caenorhabditis elegans locomotion: where do we stand? BioScience. 2014 Jun;64(6):476–86. doi: 10.1093/biosci/biu058.

34. Therrien M, Parker JA. Worming forward: amyotrophic lateral sclerosis toxicity mechanisms and genetic interactions in Caenorhabditis elegans. Front Genet. 2014;5:85. doi: 10.3389/fgene.2014.00085.

35. Nakayama J, Tan L, Li Y, Goh BC, Wang S, Makinoshima H, et al. A zebrafish embryo screen utilizing gastrulation identifies the HTR2C inhibitor pizotifen as a suppressor of EMT-mediated metastasis. Elife. 2021;10:e70151. doi: 10.7554/eLife.70151.

36. Aranda-Martínez P, Fernández-Martínez J, Ramírez-Casas Y, Guerra-Librero A, Rodríguez-Santana C, Escames G, et al. The zebrafish, an outstanding model for biomedical research in the field of melatonin and human diseases. Int J Mol Sci. 2022;23(13):7438. doi: 10.3390/ijms23137438.

37. Rouf MA, Wen L, Mahendra Y, Wang J, Zhang K, Liang S, et al. The recent advances and future perspectives of genetic compensation studies in the zebrafish model. Genes Dis. 2023 Mar;10(2):468–79. doi: 10.1016/j.gendis.2021.12.003.

38. Brown-Panton CA, Sabour S, Zoidl GSO, Zoidl C, Tabatabaei N, Zoidl GR. Gap junction Delta-2b (gjd2b/Cx35.1) depletion causes hyperopia and visual-motor deficiencies in the zebrafish. Front Cell Dev Biol. 2023;11:1150273. doi: 10.3389/fcell.2023.1150273.

39. Health Canada. Drug product database (SANDOMIGRAN) [Internet]. 2024 [cited 2025 Feb 7]. Available from: https://health-products.canada.ca/dpd-bdpp/info?code=48362&lang=eng.

40. Agence nationale de sécurité du médicament et des produits de santé (ANSM). Répertoire des Spécialités Pharmaceutiques (SANMIGRAN) [Internet]. 2024 [cited 2025 Feb 7]. Available from: http://agence-prd.ansm.sante.fr/php/ecodex/extrait.php?specid=65458611.

41. Medicines and Healthcare products Regulatory Agency (MHRA). Products website (Pizotifen) [Internet]. 2024 [cited 2024 Feb 5]. Available from: https://products.mhra.gov.uk/search/?search=pizotifen&page=1.

42. Paladin Labs. Product monograph: SANDOMIGRAN (Pizotifen hydrogen malate 0.5 mg and 1 mg tablets) [Internet]. 2012 [cited 2024 Feb 5]. Available from: http://www.paladin-labs.com/our_products/Sandomigran_en.pdf.

43. Przegaliński E, Baran L, Palider W, Siwanowicz J. The central action of pizotifen. Psychopharmacology (Berl). 1979;62(3):295–300. doi: 10.1007/BF00431961.

44. Do Nascimento BG, Rodrigues JR, Rosa Silva MC, Dutra Costa BP, Leite M, Pereira Pyterson M, et al. Serotonergic mediation of orienting and defensive responses in zebrafish. Mar Freshw Behav Physiol. 2025 Jan 10;1–16. doi: 10.1080/10236244.2025.2451826.

45. McNaught-Flores DA, Kooistra AJ, Chen YC, Arias-Montano JA, Panula P, Leurs R. Pharmacological characterization of the zebrafish (Danio rerio) histamine H1 receptor reveals the involvement of the second extracellular loop in the binding of histamine. Mol Pharmacol. 2024;105(2):84–96. doi: 10.1124/molpharm.123.000741.

46. Sarantos MR, Papanikolaou T, Ellerby LM, Hughes RE. Pizotifen activates ERK and provides neuroprotection in vitro and in vivo in models of Huntington’s disease. J Huntingtons Dis. 2012;1(2):195–210. doi: 10.3233/JHD-120033.

47. Swaminathan A, Bouffard M, Liao M, Ryan S, Callister JB, Pickering-Brown SM, et al. Expression of C9orf72-related dipeptides impairs motor function in a vertebrate model. Hum Mol Genet. 2018;27(10):1754–62. doi: 10.1093/hmg/ddy083.

48. Moreno-Mateos MA, Vejnar CE, Beaudoin JD, Fernandez JP, Mis EK, Khokha MK, et al. CRISPRscan: designing highly efficient sgRNAs for CRISPR-Cas9 targeting in vivo. Nat Methods. 2015;12(10):982–8. doi: 10.1038/nmeth.3543.

49. Samarut É, Lissouba A, Drapeau P. A simplified method for identifying early CRISPR-induced indels in zebrafish embryos using high-resolution melting analysis. BMC Genomics. 2016;17(1):547. doi: 10.1186/s12864-016-2881-1.

50. Ye J, Coulouris G, Zaretskaya I, Cutcutache I, Rozen S, Madden TL. Primer-BLAST: a tool to design target-specific primers for polymerase chain reaction. BMC Bioinformatics. 2012;13:134. doi: 10.1186/1471-2105-13-134.

51. Chuang LY, Yang CH, Lin MC. Specific primer design for the polymerase chain reaction. Biotechnol Lett. 2013;35(10):1541–9. doi: 10.1007/s10529-013-1249-8.

52. Singh J, Pan YE, Patten SA. NMJ Analyser: a novel method to quantify neuromuscular junction morphology in zebrafish. Bioinformatics. 2023;39. doi: 10.1093/bioinformatics/btad720.

53. Stiernagle T. Maintenance of C. elegans. WormBook. 2006:1–11. doi: 10.1895/wormbook.1.101.1. PMID: 18050451.

